# Towards a cumulative science of vocal markers of autism: a cross-linguistic meta-analysis-based investigation of acoustic markers in American and Danish autistic children

**DOI:** 10.1101/2021.07.13.452165

**Authors:** Riccardo Fusaroli, Ruth Grossman, Niels Bilenberg, Cathriona Cantio, Jens Richardt Møllegaard Jepsen, Ethan Weed

## Abstract

Acoustic atypicalities in speech production are argued to be potential markers of clinical features in Autism Spectrum Disorder (ASD). A recent meta-analysis highlighted shortcomings in the field, in particular small sample sizes and study heterogeneity (Fusaroli et al., 2017). We showcase a cumulative (i.e., explicitly building on previous studies both conceptually and statistically) yet self-correcting (i.e., critically assessing the impact of cumulative statistical techniques) approach to prosody in ASD to overcome these issues.

We relied on the recommendations contained in the meta-analysis to build and analyze a cross-linguistic corpus of multiple speech productions in 77 autistic and 72 neurotypical children and adolescents (>1000 recordings in Danish and US English). We used meta-analytically informed and skeptical priors, with informed priors leading to more generalizable inference. We replicated findings of a minimal cross-linguistically reliable distinctive acoustic profile for ASD (higher pitch and longer pauses) with moderate effect sizes. We identified novel reliable differences between the two groups for normalized amplitude quotient, maxima dispersion quotient, and creakiness. However, the differences were small, and there is likely no one acoustic profile characterizing all autistic individuals. We identified reliable relations of acoustic features with individual differences (age, gender), and clinical features (speech rate and ADOS sub-scores).

Besides cumulatively building our understanding of acoustic atypicalities in ASD, the study shows how to use systematic reviews and meta-analyses to guide the design and analysis of follow-up studies. We indicate future directions: larger and more diverse cross-linguistic datasets, focus on heterogeneity, self-critical cumulative approaches and open science.

**Lay Summary:** Autistic individuals are reported to speak in distinctive ways. Distinctive vocal production can affect social interactions and social development and could represent a noninvasive way to support the assessment of ASD. We systematically checked whether acoustic atypicalities highlighted in previous articles could be actually found across multiple recordings and two languages. We find a minimal acoustic profile of ASD: higher pitch, longer pauses, increased hoarseness and creakiness of the voice. However, there is much individual variability (by age, sex, language, and clinical characteristics). This suggests that the search for one common “autistic voice” might be naive and more fine-grained approaches are needed.

## 1. Introduction

Atypical prosody and voice are commonly-reported aspects of the speech of people with autism, which has been characterized as flat, sing-songy, pedantic, hollow, inappropriate, hoarse or hypernasal (Asperger, 1991; Baltaxe & Simmons, 1975; Goldfarb et al., 1956; Kanner, 1943; Pronovost et al., 1966). Indeed, distinctive prosody is part of the diagnostic criteria in the ICD-10 and in the ADOS-2 assessment for autism (Lord et al., 2009; World Health Organization, 1992) and is indicated as one of the earliest-appearing markers of a possible Autism Spectrum Disorder (ASD) diagnosis (Oller et al., 2010). These vocal factors may play a role in the socio-communicative impairments associated with the disorder. In addition to potentially impeding effective communication of e.g., emotional content (Travis & Sigman, 1998), they also generate negative responses from neurotypical raters, even when hearing as little as 1 second of speech (Grossman, 2015; Sasson et al., 2017). These negative first impressions may have long term effects, e.g., providing a less optimal scaffolding for socio-communicative development, or even increasing the risks of social withdrawal and anxiety (Fay & Schuler, 1980; Fusaroli et al., 2019, 2021; Paul et al., 2005; Shriberg et al., 2001; Van Bourgondien & Woods, 1992; Warlaumont et al., 2014). Given their potential role in affecting social functioning and in assisting diagnostic and assessment processes, it is important to understand how these vocal atypicalities manifest themselves across autistic people and uncover their acoustic underpinnings. This is especially true if we want to assess whether and how assessment and intervention tools should be developed to target them.

It has been nearly 80 years since unusual prosody was first reported by Kanner in 1943 (Kanner, 1943), and there is a growing interest in finding markers of ASD and social functioning. Nevertheless, two reviews of the field show that we know remarkably little about the precise perceptual and acoustic properties differentiating the speech of autistic people from that of neurotypical peers. A review of the literature from 2003 concluded that “No study offers a large number of subjects, matched with neurotypical children or adults (controlled for linguistic and non-verbal abilities). If findings were consistent, small-scale studies would offer pointers, but as it is these do not inspire confidence” (McCann & Peppé, 2003, p. 347). A more recent systematic review and meta-analysis (Fusaroli et al., 2017) concluded that single acoustic features (pitch mean and variability^1^) showed robust small to moderate differences between groups. However, the studies reviewed were noted to have small sample size, have high heterogeneity in methods and features analyzed, largely neglect voice quality features (which are highlighted as important by speech pathologists and speech processing research), and leave a lingering need for multivariate approaches to account for shared variance and interactions across features. In other words, there is a need for a more rigorously cumulative and yet self-correcting scientific approach to the understanding of vocal and prosodic atypicalities in ASD. We define as “cumulative” an approach that *explicitly and systematically builds upon previous studies in terms of both study design and statistical inference*. We define as self-correcting an approach that *does not take previous findings at face value, but explicitly assesses them in the light of new studies and provides critical input for future research* (e.g., by assessing the difference that including previous findings in the current analysis makes for the statistical inference).

In this paper we develop such an approach. We rely on the most recent systematic review and meta-analysis of the field to set up the analysis of two new datasets. We build on the recommendations there produced, and test the replicability of updated meta-analytic results (Fusaroli et al., 2018).

### 1.1. Towards a cumulative research approach

A very common approach to cumulative research is to perform systematic reviews to map the field, and meta-analyses of previous results to achieve a more robust estimate of the underlying phenomena, beyond the variability of single studies. As an example, of the 17 studies conducted between 2010 and 2016, 13 found that autistic people had a wider pitch range, while 4 studies found the opposite effect (Fusaroli et al., 2017). A meta-analysis can pool the data from the different studies and perform an overarching inference as to the underlying effect size, and even assess whether systematic variations in study design (e.g., monological vs. dialogic speech production) might explain the differences in effects between studies (Cox et al., 2021; Cumming, 2014; Nguyen et al., 2021; Parola et al., 2020; Weed & Fusaroli, 2020). A common critique of this approach is “garbage-in-garbage-out”: If the studies included are too diverse, biased, or methodologically problematic, the meta-analytic inference will also be unreliable, and potentially overestimate effect sizes (Lewis et al., 2020; Open Science Collaboration, 2015). While a few different techniques have been developed to assess the heterogeneity between studies and potential publication biases (Dwan et al., 2013), they are not a solution to the issue of more reliably estimating the true effect, and the critique remains valid (Rocca & Yarkoni, 2021). Systematic reviews and meta-analyses are invaluable to get a feel for the field and identify potential issues or directions for research, but they should always be taken with caution as the researchers have no control on the quality and biases of the studies reviewed. We therefore need to critically combine systematic assessments of the field with well-targeted replications and new studies.

#### 1.1.1 Building on existing guidelines

Previous systematic reviews and meta-analyses can be used to identify current best practices, pitfalls and blindspots, and therefore develop *guidelines for new studies* (Gelman et al., 2008; König & van de Schoot, 2018; Williams et al., 2018). Indeed, (Fusaroli et al., 2017) identified several key areas for improvement in investigating vocal atypicalities in ASD.

##### 1.1.1.1 More attention to the heterogeneity of the disorder

Building on insights from Fusaroli et al (2017), we designed a new study based on two existing corpora of voice data, collected in the US and Denmark (Cantio et al., 2016; Grossman et al., 2013). The study involves a high degree of heterogeneity in its sample: two diverse languages (Danish and US English) and a larger than average sample: 77 autistic participants and 72 neurotypical (NT) participants, against a previous median sample size of 17. Further, the study involves repeated measures of voice (between 4 and 12 separate recordings per participant). For each participant we have demographic (age, biological sex, native language) and clinical features (ADOS total scores, as well as the following sub-scores: Communication, Social Interaction, and Restricted and Repetitive Behaviors).

##### 1.1.1.2. More systematic use of acoustic features across studies

Second, (Fusaroli et al., 2017) noted that different studies measured different acoustic features with diverse methods, without any explicit concern about comparing across studies^2^. Within our sample we systematically extract the acoustic features identified in the updated meta-analysis by (Fusaroli et al., 2018). This includes measures of pitch (median and variability), and rhythm (speech rate, average syllable length, pause number per unit of time, and pause length)^3^.

Further, clinicians variously describe autistic voices as hoarse, creaky, breathy, harsh or otherwise dysphonic (Baltaxe, 1981; Pronovost et al., 1966; Sheinkopf et al., 2000). We therefore identified in the speech signal processing literature acoustic features thought to be related to these perceptual qualities, e.g., pertaining to the glottal or spectral domain, fully listed in the methods section, and in Table S1.

Of course, by expanding the acoustic features investigated we risk producing a non-trivial increase in the complexity of the research, and in the number of statistical analyses required. Further, acoustic features are likely to be related to each other, and therefore we should assess whether all the features investigated provide independent information, and whether it is really necessary to add more complex acoustic measures of voice quality to the more traditional prosodic measures. Accordingly, we provide several analyses in the appendix including principal component and network analyses exploring the variance shared across features (Appendices S4 and S7)

##### 1.1.1.4. Open science practices

To further promote cumulative approaches, we also provide an open de-identified dataset including demographic, clinical and acoustic features, and open scripts to reproduce our analysis on the current dataset and replicate and extend our findings on future datasets (https://osf.io/gnhw4/?view_only=3e51ee6253d548eb836af23ed9d01d73m; see also Parish-Morris et al., 2016; Wilkinson et al., 2016). Note that the original speech recordings cannot be shared as they are considered identifiable data. We therefore – in line with our consent forms and current data privacy regulations - only share the acoustic features as employed in the analyses (summary statistics at the recording and 6-second segments levels).

### 1.2. Hypotheses

Based on the systematic review and meta-analysis and on current meta-scientific knowledge on replicability of meta-analytic findings, we developed the following expectations.

1. We will replicate meta-analytic findings that autistic people have: higher pitch mean and variability; more frequent and longer pauses; no differences in speech rate and syllable length, compared to neurotypical participants.

a. Effect sizes will be half to a third smaller than previous meta-analytic findings due to hard to correct publication bias issues (Kvarven et al., 2020; Lewis et al., 2020). See Table 2 for effect sizes of meta-analytic findings.
2. At least some of the measures of voice quality will be different in autistic people compared to neurotypicals, with effect sizes comparable to prosodic measures.
3. We expect the acoustic profile of autistic voice to be affected by individual differences (vs. a unique profile of autistic voice). In particular, we expect effects to be different by gender, and age. For instance, a mega-analytic study (Fusaroli et al. 2018) found that the acoustic markers of ASD were particularly pronounced in older individuals and in the more numerous male group· We also expect acoustic features to relate to clinical features of ASD as measured by ADOS sub-scores. In particular, increased pitch mean and variability, and pause number will relate to increased sub-scores, plausibly Social and Communication.

**Table 1.**
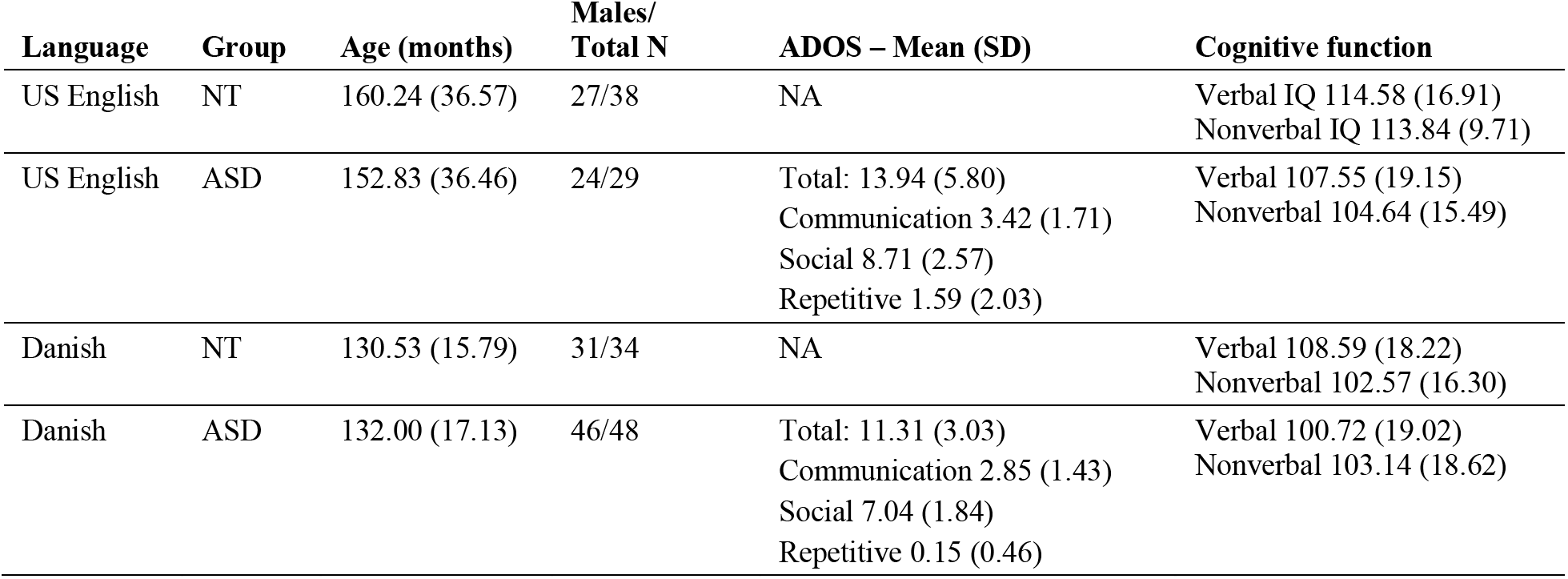
Participant characteristics. Clinical symptoms severity was measures using the Autism Diagnostic Observation Schedule – Generic (Lord et al., 2008). Cognitive functions were measured using the WISC-III for the Danish data (Kaufman, 1994), and the Leiter-R (nonverbal IQ, Roid & Miller, 1997) and the Peabody Picture Vocabulary test (receptive vocabulary, Dunn & Dunn, 2007).

| **Language** | **Group** | **Age (months)** | **Males/**<br>**Total N** | **ADOS – Mean (SD)** | **Cognitive function** |
| --- | --- | --- | --- | --- | --- |
| US English | NT | 160.24 (36.57) | 27/38 | NA | Verbal IQ 114.58 (16.91)<br>Nonverbal IQ 113.84 (9.71) |
| US English | ASD | 152.83 (36.46) | 24/29 | Total: 13.94 (5.80)<br>Communication 3.42 (1.71)<br>Social 8.71 (2.57)<br>Repetitive 1.59 (2.03) | Verbal 107.55 (19.15)<br>Nonverbal 104.64 (15.49) |
| Danish | NT | 130.53 (15.79) | 31/34 | NA | Verbal 108.59 (18.22)<br>Nonverbal 102.57 (16.30) |
| Danish | ASD | 132.00 (17.13) | 46/48 | Total: 11.31 (3.03)<br>Communication 2.85 (1.43)<br>Social 7.04 (1.84)<br>Repetitive 0.15 (0.46) | Verbal 100.72 (19.02)<br>Nonverbal 103.14 (18.62) |

**Table 2.** Estimated standardized mean differences (ASD – NT) for the six acoustic measures present in the meta-analysis, as estimated separately by the meta-analytically informed and the skeptical models. The first column reports the main effect of the diagnostic group (across sex and age), respectively from the meta-analysis (MA), for the skeptical Danish and US English models, and for the informed ones. The second column indicates the interaction between the effect of the diagnostic group and biological sex (Male – Female), that is, the difference in effect of group between the male and the female participants. The third column reports the interaction between the effect of diagnostic group and age, that is, the change in effect size as age increases by 1 standard deviation. ER indicates the evidence ratio for the difference, ER01 the evidence ratio for the null effect. Bold text indicates findings for which there is more than anecdotal evidence (Evidence Ratio above 3).

|  | Group (ASD - NT) | Group *<br>Biological sex (M - F) | Group *<br>Age |
| --- | --- | --- | --- |
| *Pitch Median* | Informed Model Weight = 0.79 |  |  |
| MA | 0.38 (0.16 0.59) | NA | NA |
| Skeptical DK | **0.12 (-0.08 0.32) ER = 5.26** | **0.32 (-0.02 0.64) ER = 14.62** | -0.01 (-0.06 0.03) ER = 2.34<br>ER01 = 9.73 |
| Skeptical US | **0.36 (0.12 0.61) ER = 221** | -0.07 (-0.46 0.33) ER = 1.56<br>ER01 = 1.81 | **-0.02 (-0.06 0.02) ER = 4.13** |
| Informed DK | **0.27 (0.14 0.41) ER > 1000** | NA | NA |
| Informed US | **0.43 (0.28 0.59) ER > 1000** | NA | NA |
| *Pitch Variability* | Informed Model Weight = 0.97 |  |  |
| MA | 0.48 (0.26 0.7) | NA | NA |
| Skeptical DK | **0.31 (0.16 0.46) ER > 1000** | **0.28 (-0.06 0.61) ER = 11.05** | **0.01 (-0.04 0.05) ER = 1.45**<br>**ER01 = 10.29** |
| Skeptical US | 0.02 (-0.2 0.23) ER = 1.3 ER01 = 2.32 | **-0.23 (-0.66 0.2) ER = 4.41** | **-0.02 (-0.06 0.01) ER = 6.02** |
| Informed DK | **0.41 (0.29 0.52) ER > 1000** | NA | NA |
| Informed US | **0.28 (0.14 0.43) ER > 1000** | NA | NA |
| *Speech Rate* | Informed Model Weight = 1 |  |  |
| MA | 0.02 (-0.27 0.31) | NA | NA |
| Skeptical DK | 0.03 (-0.14 0.2) ER = 1.55 ER01 = 2.71 | -0.02 (-0.35 0.29) ER = 1.19<br>ER01 = 2.15 | 0 (-0.04 0.04) ER = 1.12 ER01 = 11.28 |
| Skeptical US | **-0.11 (-0.28 0.05) ER = 6.74** | **0.24 (-0.21 0.69) ER = 4.12** | **-0.02 (-0.06 0.01) ER = 8.71** |
| Informed DK | 0.03 (-0.11 0.17) ER = 1.75<br>ER01 = 1.62 | NA | NA |
| Informed US | **-0.09 (-0.22 0.06) ER = 5.06** | NA | NA |
| *Syllable Length* | Informed Model Weight = 0.98 |  |  |
| MA | 0.06 (-0.63 0.76) | NA | NA |
| Skeptical DK | -0.02 (-0.14 0.09) ER = 1.64<br>ER01 = 4.19 | -0.03 (-0.3 0.25) ER = 1.37<br>ER01 = 2.38 | 0 (-0.04 0.04) ER = 1.14 ER01 =<br>13.57 |
| Skeptical US | 0 (-0.21 0.22) ER = 1 ER01 =<br>2.16 | -0.04 (-0.52 0.45) ER = 1.28<br>ER01 = 1.48 | 0 (-0.03 0.04) ER = 1.1 ER01 =<br>13.81 |
| Informed DK | -0.02 (-0.13 0.09) ER = 1.6 ER01<br>= 5.01 | NA | NA |
| Informed US | 0.01 (-0.22 0.23) ER = 1.07<br>ER01 = 2.58 | NA | NA |
| *Pause Number* | Informed Model Weight = 1 |  |  |
| MA | 0.4 (0.01 0.78) | NA | NA |
| Skeptical DK | **-0.11 (-0.24 0.01) ER = 14.04** | -0.05 (-0.34 0.22) ER = 1.6<br>ER01 = 2.55 | -0.01 (-0.05 0.03) ER = 2.36<br>ER01 = 11.55 |
| Skeptical US | **-0.17 (-0.35 0) ER = 17.6** | **0.39 (-0.05 0.83) ER = 12** | **-0.02 (-0.05 0.01) ER = 4.66** |
| Informed DK | -0.04 (-0.17 0.08) ER = 2.41<br>ER01 = 15.95 | NA | NA |
| Informed US | **-0.09 (-0.26 0.09) ER = 4.18** | NA | NA |
| *Pause Length* | Informed Model Weight = 1 |  |  |
| MA | 0.21 (-0.09 0.5) | NA | NA |
| Skeptical DK | **0.21 (0.03 0.39) ER = 34.09** | **0.15 (-0.19 0.47) ER = 3.4** | 0.01 (-0.04 0.05) ER = 2.05<br>ER01 = 10.31 |
| Skeptical US | **0.27 (0.11 0.43) ER = 332** | -0.12 (-0.55 0.31) ER = 1.96<br>ER01 = 1.4 | 0 (-0.03 0.03) ER = 1.17 ER01 =<br>16.13 |
| Informed DK | **0.24 (0.08 0.39) ER = 153** | NA | NA |
| Informed US | **0.26 (0.12 0.4) ER = 999** | NA | NA |

In a more exploratory fashion, we comparatively assess the use of meta-analytically informed and skeptical priors on model quality (assessed via model comparison) and estimates.

## 2. Materials and Methods

### 2.1. Participants and recordings

We collected two Danish and US English datasets involving 77 autistic participants and 72 neurotypical (NT) participants, all with verbal and non-verbal cognitive function within a typical range (see Table 1 for details). Each participant recorded several audios, for a total of 1074 unique recordings. The Danish dataset included 29 autistic participants (335 recordings) and 38 NT participants (427 recordings), retelling stories (Memory for stories, Reynolds & Voress, 2007) and freely describing short videos (Abell et al., 2000). The US English dataset included 48 autistic (178 recordings) and 34 NT (134 recordings) participants, retelling stories (Grossman et al., 2013). The recordings had been collected for other purposes and their content – but not acoustics – analyzed in published studies (Cantio et al., 2016; Grossman et al., 2013).

Note that the two samples are only roughly matched. While cognitive function and clinical features as measured by ADOS are largely overlapping, US participants are a bit older than Danish ones, present a larger variability in age and a slightly higher number of female participants. Further, the language spoken, while in both cases a Germanic one, is obviously different. In particular, Danish is often characterized as an atypical language with strong reduction in consonant pronunciation (Trecca et al., 2021), although no systematic comparison with US English has been performed to our knowledge. These differences are not an issue for the following analyses, given that the effects are tested separately in the two corpora. It is crucial to assess whether so called vocal markers of ASD can generalize across corpora with different characteristics and explore how these differences might matter for the generalizability of the findings.

Note also that the number of autistic girls in the sample is limited: 7 in total, in line with the higher detection of autism in males. Therefore, the estimates are to be taken with much caution. Since future studies might include our estimates in a cumulative approach, we nevertheless report them.

All recordings were pre-processed to remove background noise and interviewer speech when present. 32 acoustic measures were extracted (see Table 2 and 3). A full description of the process and features is available in the Supplementary Materials – S1.

**Table 3.**
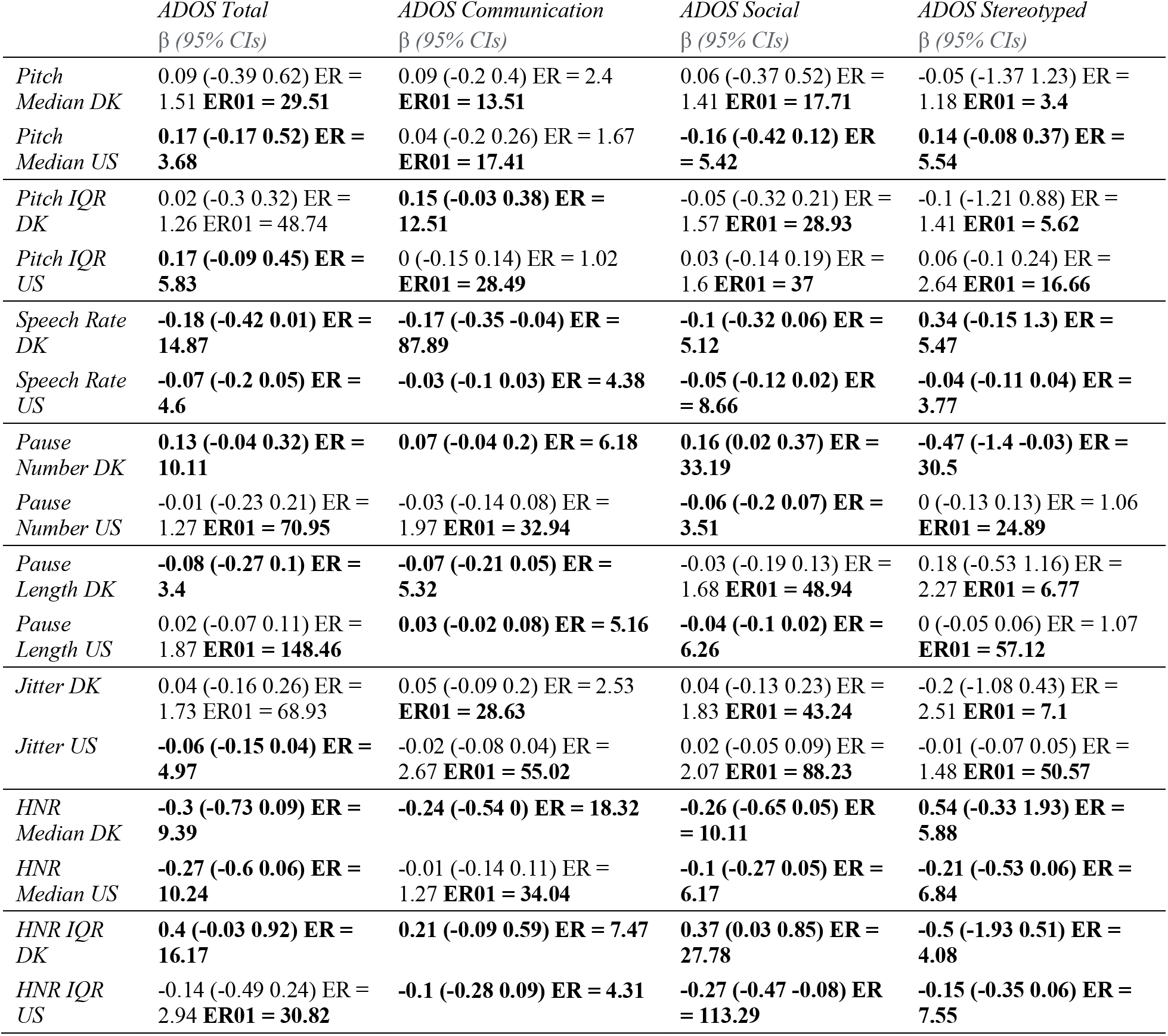
Estimated standardized relation between acoustic and clinical features. ER indicates the evidence ratio for the difference, ER01 the evidence ratio for the null effect.

### 2.2. Statistical modeling

#### 2.2.1 Updated meta-analytic estimates

Effect size estimates from previous studies were retrieved from metavoice.au.dk, a community augmented meta-analysis website, which integrated and updated the estimates collected by (Fusaroli et al., 2017, 2018). We re-ran the meta-analysis on this updated dataset in accordance with the original specifications. In other words, we used a multilevel model with standardized effect sizes (Cohen’s d) as outcome and accounting for varying effects by article and sample (some articles re-used the same participants). This yielded meta-analytic effect sizes (including measures of uncertainty) for pitch mean and variability, average speech rate, syllable and pause average duration and number of pauses per second. The estimates are presented in Table 2 and the script is provided on the OSF repository (https://osf.io/gnhw4/?view_only=3e51ee6253d548eb836af23ed9d01d73s).

#### 2.2.1. Differences by diagnostic group

For ease of comparison with the standardized effect sizes used in the meta-analysis, we standardized our acoustic features, that is, we centered them on the mean, and divided them by the standard deviation.^4^ To assess whether autistic participants differed from NT participants, we ran Bayesian multilevel Gaussian regression models with the standardized acoustic feature as outcome, group (ASD vs. NT) and language (Danish vs. US English) as predictors (separately assessing the effects within language), and varying effects by participant (separately by language and group). We contrasted the results achieved with meta-analytically informed priors - when available - and weakly skeptical priors - that is, with expectations of no or small effects, thereby conservatively regularizing the inference and reducing overfitting. We were interested in both whether the effects would be robust to the change of priors, and whether the informed or the skeptical priors would lead to more robust inference, that is, would lead to lower estimated out-of-sample error. To assess the import of using meta-analytic and skeptical priors, we report the same model estimates for both models. Further, we adopted a Leave-One-Out model comparison framework estimating the model’s out-of-sample performance, in other words, estimating the ability of the model to generalize to new data (Yao et al., 2018). We then calculated their relative stacking weight based on Leave-One-Out Information Criteria, assessing the probability of each model to be better than the other on a 0-1 scale. When one model gets a stacking weight of 1, that model is reliably better than the other. When the scores are closer to 0.5, both models provide valuable information in predicting new data and should be both considered for future work. This procedure informs us as to whether adding information from previous findings in our statistical models enabled us to create more robust models, with greater estimated chances of our findings replicating in new studies. We reported the estimated difference by group in terms of mean difference separately by language (that is, by corpus), 95% Compatibility Intervals (CIs, indicating the probable range of difference, assuming the model is correct and the data representative of the population) and Evidence Ratio (ER, evidence in favor of the effect observed against alternative hypotheses). When ER was weak (below 3, that is, less than three times as much evidence for the effect as for alternative hypotheses), we also calculated the Evidence Ratio in favor of the null hypothesis. Note that given the standardization of the outcome variables, the effect size is equivalent to Cohen’s d, that is, is expressed in units of standard deviations.

To evaluate the potential role of individual differences in biological sex (Male vs. Female) and age, we built additional models, one per each suggested moderator interacting with group separately in the two languages. Age was modeled in terms of years and scaled. We reported the model estimates for the interaction, including CIs and ERs.

Further details on the implementation and on the priors used are presented in the Supplementary Materials – S2, and S5. Note also that we report additional analyses in the appendix to assess the robustness of the findings: we repeat all analyses on audio segments of 6 seconds to control for recoding length, see Tables S4, S5, S6. The results generally support our main findings and we report in the manuscript only qualitative divergences.

#### 2.2.2. Relations to clinical features

To analyze the relation of acoustic and clinical features (ADOS total, Communication, Social Interaction, Repetitive Behaviors scores) we built multilevel Bayesian linear regression models with the acoustic feature as outcome (rescaled on a 0-1 scale) and clinical features as ordinal predictors, on the ASD group only, separately by language and with varying effects by participant (separately by language). We selected only features that were highlighted by the metaanalysis, as associated with group differences (pitch median and variability, speech rate, pause number and length), or with clinical features (jitter, Harmonic to Noise Ratio).

We otherwise followed the procedure described in the previous paragraphs. Further details on the implementation and priors are available in the Supplementary Materials – S3 and S5. Note that given the rescaling of the outcome and predictor variables, the effect size is on the scale of Pearson’s r.

The data analysis scripts are available in the article repository at Open Science Foundation (https://osf.io/gnhw4/?view_only=3e51ee6253d548eb836af23ed9d01d73), and further details on the software employed is available in the Supplementary Materials – S5.

## 3. Results

### 3.1. Analysis of group differences in acoustic features

#### 3.1.1. Acoustic features with meta-analytic results

The detailed results and comparison to the meta-analysis are reported in Table 2, and Figure 1. Using informed meta-analytic priors yielded bigger differences than using skeptical ones, however they were still smaller than in the meta-analysis. Interestingly all informed models performed better than the skeptical ones. In other words, including information from previous studies often made our statistical inference more robust and able to generalize to new data (LOO based stacking weights for informed models always above .75). This suggests that a more consistent practice of “posterior passing” (Brand et al., 2019), that is, of using previous findings as priors in current studies, would lead to more robust inferences. The results generally supported our broader hypotheses, if not the more specific details. We mostly replicated meta-analytic findings across both datasets (H1). Autistic participants across languages tend to use higher pitch, as well as fewer and longer pauses, and showed no differences in syllable length. Perhaps unsurprisingly, the effect sizes in our data are often smaller than in the meta-analysis (H1a), except for length of pauses. We also observe evidence for the importance of individual and linguistic differences (H3). Only in US English did we see robust evidence of slower speech rate and only in Danish did we see increased pitch variability. While previous studies (Fusaroli et al., 2018) suggested that vocal markers of ASD would be stronger in younger boys, we observe a more complex picture: biological sex and age interact with the effects, but inconsistently so across languages. Further given the small number of female participants involved, much caution should be exercised.

The findings are maintained if using only 6 second clips of the audio recordings, with the exception of syllable length becoming credibly longer in ASD in both languages and the advantage of using meta-analytic priors being reduced (see Supplementary Materials – S9, and Figure S3).

#### 3.1.2. Novel acoustic features

The detailed results for each of the 26 features are reported in Table S2 in the appendix. We observe small to moderate (< 0.4) but reliable differences by group in the voice quality features within each dataset, which are comparable to those in prosodic features (partially corroborating H2). As in more traditional acoustic features we see that including biological sex and age of the participants does in some cases affect the group differences (corroborating H3). However, strikingly, only three acoustic measures present the same small but reliable difference between the diagnostic group across the two languages (questioning the generalizability of H2). In particular, autistic participants have higher Normalized Amplitude Quotient (NAQ, effect sizes of 0.1 and 0.11), Maxima Dispersion Quotient (MDQ, effect sizes of 0.07 and 0.06) and creak (effect sizes of 0.13 and 0.15).

### 3.2. Relation with clinical features

Detailed results are presented in Table 3. While we can observe several reliable relations between acoustic and clinical features, the only consistent one across languages is speech rate (the slower the speech, the more severe the clinical feature), and to a lesser degree Harmonic to Noise Ratio (the lower, the more severe the clinical feature). Many correlations are small (< 0.2 or 4% of the variance), but some are moderate (between 0.4 and 0.54, that is, between 16% and 29% of the variance). The findings are analogous, albeit with smaller effect size in the 6 second audio recordings (see Supplementary Materials – S9).

## 4. Discussion

In this work we aimed at building the foundations for a cumulative yet self-correcting approach to the study of prosody in ASD. Relying on a previous systematic review and an updated metaanalysis of the field, we hypothesized that: H1) meta-analytic findings would replicate, potentially with smaller effect sizes; H2) voice quality measures would yield differences in the 2 groups analogous in size to those from prosodic measures; H3) individual demographic, clinical and linguistic differences would play an important role, defying the idea of a unique acoustic profile of ASD. We also assessed whether the use of informed priors would improve the generalizability of the statistical findings, and – in the appendix - whether the acoustic features showed obvious redundancies, allowing to reduce their numbers. In the following discussion we will consider how the findings bear on the hypotheses and explorations, highlight the limitations of the current study, and discuss further the cumulative yet self-correcting approach we propose.

Traditionally, given a previous finding, a replication of that finding is the production of similar results (same direction of effects and comparable effect sizes) in a new study following analogous experimental and/or statistical procedures (Goodman et al., 2016), while not finding the same pattern of results indicates a failed replication. Further, if the replication attempts to apply the results of a study “to populations with for instance a different language, age distribution, or other demographic and clinical characteristics” (Vandenbroucke et al., 2007), we call this a generalization. Most previous findings and current results did not replicate in a generalizable fashion. We found a minimal (characterized by only few features) acoustic profile of ASD across Danish and US English: Autistic participants tended to use higher pitch, fewer but longer pauses, and increased NAQ, MDQ and creak compared to NT participants. Given the heterogeneity of previous studies and uncertainty about publication bias reported in the meta-analysis, even these minimal cross-linguistically generalized replications are far from trivial. However, equally important is the focus that our findings put on linguistic, demographic, and clinical differences undermining the idea of a strong acoustic profile of ASD. There are many language-specific effects (e.g., speech rate being slower in ASD only for US English), and demographic differences (sex and age) affect even the cross-linguistically reliable acoustic features of ASD, albeit not in the same directions across languages and acoustic features. Additionally, there were intriguing moderate relations between acoustic measures and clinical symptoms. However, only speech rate and – to a lower extent - HNR show the same cross-linguistic relation: the slower the speech, and the lower the HNR, the more severe the clinical feature. Even more tellingly, these features are not consistently different between diagnostic groups.

The findings thus do not lead to sweeping statements on vocal markers of ASD and the role of age- and sex-related differences, except that there might be no one general extensive acoustic profile of ASD. Systematic individual variations (sex, age, language, clinical features) should be always taken into account and we suspect that multiple clusters of acoustic profiles in ASD could be identified, all leading to the more general clinical descriptions of vocal atypicalities in ASD. However, to explore this idea and its potential clinical applications, we need to construct even larger cross-linguistic datasets systematically covering the heterogeneity in clinical features - and beyond - of autistic people, and more explicit normative modeling of individual variability (Marquand et al., 2019).

Our exploration of feature reduction methods (appendix S4 & S7) did not yield any clear finding. Future directions should explicitly include machine learning techniques targeting diagnostic group differences and relevant clinical features (Rybner et al., 2021).

More generally our study provides a concrete case of cumulative yet self-critical scientific approach, which might extend beyond the specific phenomenon investigated. The need to systematically rely on previous literature (e.g., via systematic reviews and meta-analyses) has been argued to stand in contrast with the potential unreliability of the current literature due to questionable research practices, publication bias and other issues (Bavel et al., 2016; Benjamin et al., 2018; Fabrigar et al., 2020; Kenny & Judd, 2019; Kvarven et al., 2020; Loken & Gelman, 2017; Maxwell et al., 2015; Oberauer & Lewandowsky, 2019; Yarkoni, 2020). In this study, we suggest a way to account for both sides. We rely on a previous systematic review to design the current study, and on the (updated) meta-analytic findings to set up priors for our analyses. At the same time, we critically attempt to replicate previous findings and compare statistical inferences relying on meta-analytical priors with inferences relying on skeptical priors. Estimations of out-of-sample errors indicate that including the meta-analytic findings in our analyses actually improves the generalizability of our inferences. This direct way of producing cumulative results via “posterior passing” (Brand et al., 2019) thus seems quite promising. There are of course some cautions to consider. The first is that estimated out-of-sample errors assume that new recordings would belong to the same population described by the current sample of recordings. This means that it is not a reliable estimate of actual error when assessing e.g., new recordings in a different language or a population with different clinical profiles. Further, our results indicate that the advantage of using meta-analytic priors is not as clear when analyzing more controlled samples (clips of 6 seconds analyzed in the Appendix). This suggests that the usefulness of posterior passing, especially when the meta-analysis suggests the presence of noise and heterogeneity, might be lessened in the presence of more controlled measurements, and possibly larger datasets. Nevertheless, even in these conditions, we argue that comparatively applying meta-analytically informed and skeptical priors might provide a check of the inferential robustness (do the results agree across the two models?) and a measure of how the current study relates to the previous state of the art of the literature (does it fall within the range of the previous studies, or does it suggest that our current study is saying something different?).

## 5. Conclusions

We set out to more cumulatively advance the study of acoustic markers in ASD, applying and assessing the recommendations and findings in a recent systematic review and meta-analysis. Across a relatively large cross-linguistic corpus, we identified a minimal acoustic profile of ASD (higher pitch, fewer and longer pauses, higher NAQ, MDQ and creak). However, we also highlight that individual differences in language, sex, age and clinical features relate to systematic variations in the acoustic properties of speech. This suggests that the search for a population-level marker might be naive and more fine-grained approaches are needed. We released the data and scripts used in the article to facilitate such future cumulative advances. The current study critically showcases a cumulative yet self-correcting approach, which we advocate should be more commonly used.

## Acknowledgments

We are very grateful to the participants in our studies as well as to the researchers and clinical practitioners who in various guises supported the collection of the data we analyzed in this project. We wish to thank our funding sources: the Interacting Minds Centre seed fundings “Clinical Voices” and “Clinical voices in the wild”; and the Danish Independent Research Council collective project “The Puzzle of Danish”. Riccardo Fusaroli has been a consultant for F. Hoffmann-La Roche on related but not overlapping topics. The other authors have no real or potential conflicts of interest that could have had influenced the research.

## Footnotes

1 Pitch is defined the fundamental frequency of the voice transformed on a log scale to better match how it is perceived by human listeners.

2 Exploration is a necessary component of research, and one should not put standardization in front of it, to avoid getting stuck with suboptimal methods (Devezer et al., 2019; Würbel, 2000). However, it is just as important, especially when discussing markers of disorders, to assess whether the findings generalize to new samples and how different methods compare to each other (Rocca & Yarkoni, 2021).

3 Speech rate is defined as the average amount of syllables per unit of time, usually per minute, calculated including also time without actual speech, e.g., pauses. Syllable length as the average duration of a syllable, calculated dividing the total amount of spoken time by the total amount of syllables uttered. Pauses are defined as segments of recording without speech in them, usually lasting more than 200 ms (or other thresholds). Note that we did not include intensity-based measures, because we deemed them unreliable, due to their strong dependence on distance from the microphone, movements, etc. (Barsties & De Bodt, 2015).

4 Some features could have been modeled following non-Gaussian distributions (e.g., lognormal for pause length). However, keeping a Gaussian likelihood function enables an easier comparison with the meta-analytic results, as previous studies assumed Gaussian distribution of errors.

