## Supplementary Materials for "Towards a cumulative science of vocal markers of autism: a cross-linguistic meta-analysis-based investigation of acoustic markers in American and Danish autistic children"

The supplementary materials contain additional information about:

- S1 – Audio Processing and feature extraction
- S2 – Informed and skeptical priors for differences by diagnostic group (methods)
- S3 – Relations with clinical features (methods)
- S4 – Interdependence between features and feature selection (methods)
- S5 – Software implementation notes
- S6 - Differences by diagnostic group – novel features (results)
- S7 – Dimensionality reduction
- S8 – Results using 6 second snippets
- S9 – Machine learning approaches
- S10 – Steps towards a cumulative yet self-critical approach

### S1. Audio preprocessing and feature extraction.

All audio recordings were carefully listened to identify potential audio quality issues that would lead to exclusion of the recording. All audio recordings were then manually timecoded, to identify the segments corresponding to the participants’ story telling or video description, thus excluding instructions about the experiments, questions, backchannels from the interviewer (such as ‘hmmm’, or ‘ok’), etc. Once identified the relevant audio segments, we proceeded to clean the audio. Background room noise, reverberation and hum were removed from the audio recordings using iZotope RX 6 Elements^TM^. Long-term average spectra for each recording were inspected for possible noise artefacts and further cleaned if any were found.

We first extracted measures of rhythm and duration for the full relevant segment (e.g., one video description) using the open-source script syllablev2 for Praat (Boersma & Weenink, 2021; de Jong & Wempe, 2009). This generated 4 measures: speech rate (number of syllables per second, including pauses), average syllable length (excluding pauses), average number of pauses per second, and average pause length.

We extracted 15 pitch and voice quality measures every 10 milliseconds using Covarep for Matlab (Degottex et al., 2014). The measures included both spectral and glottal properties of voice: fundamental frequency (pitch), jitter, shimmer, normalized amplitude quotient (NAQ), quasi-open-quotient (QOQ), relative amplitude (H1H2), parabolic spectral parameter (PSP), harmonic richness factor (HRF), maxima dispersion quotient (MDQ), cepstral peak slope (CPS) and prominence (CPP), shape parameter of the Liljencrants-Fant Model of the Glottal Pulse Dynamic (RD), harmonic to noise ratio (HNR), creak, and clarity of articulation (see Table S1 for a description). Median and interquartile range were calculated for each of these measures using R (R Core Team, 2018). This yielded 28 measures of pitch and voice quality, for a total of 32 including those of rhythm.

The choice of pitch and rhythm measures was motivated by their widespread use in the study of vocal markers of ASD (Fusaroli et al., 2017). The choice of voice quality features was motivated by an informal review of the speech processing literature (e.g., for an overview, Cummins et al., 2015). We opted to use median and interquartile range of acoustic measures, contrary to more commonly used mean, standard deviation and range, because they are more robust to measurement errors (e.g., erroneous jumps of octave in the pitch detection algorithm). Further, fundamental frequency and other acoustic features of the human voice often present long tailed distributions, in which case mean and standard deviation become strongly correlated and determined by the tail. Median and interquartile range more robustly present independent measures respectively of mode and variance of the distribution.

*Table S1 – Description of the voice quality features extracted from the recordings.*

| Feature | Description | Reference |
| --- | --- | --- |
| Jitter | Small cycle-to-cycle variations in respectively the glottal pulse timing in voiced regions. Associated with creak and disphonicity. | (Tsanas et al., 2012) |
| Shimmer | Small cycle-to-cycle variations in respectively the glottal pulse amplitude in voiced regions. Associated with creak and disphonicity. | (Tsanas et al., 2012) |
| Harmonics to Noise Ratio (HNR) | Ratio of harmonics to inharmonic (spectral components which are not a whole number multiple of F0) components. Associated with hoarseness. | (Hanson, 1995) |
| Relative amplitude (H1H2) | Amplitude of the first harmonic relative to second harmonic. Associated with breathiness. | (Henrich et al., 2001) |
| Normalized Amplitude Quotient (NAQ) | Parametrization of the glottal closing phase using two amplitude-domain measurements from waveforms estimated by inverse filtering. Associated with breathiness. | (Alku et al., 2002) |
| Quasi-Open-Quotient (QOQ) | Relative open phase duration of a glottal cycle. Associated with breathiness. | (Szkiełkowska et al., 2018) |
| Maxima Dispersion Quotient (MDQ) | Sharpness of the glottal excitation. Associated with tense-lax properties of the voice and relatedly with breathiness. | (Gobl et al., 2015) |
| Parabolic Spectral Parameter (PSP) | A quantification of the glottal volume velocity waveform. Associated with breathiness and tense phonation. | (Alku et al., 1997) |
| Harmonic Richness Factor (HRF) | Ratio between the sum of the amplitudes of harmonics, and the amplitude at the fundamental frequency, quantifying the amount of harmonics in the magnitude spectrum of the glottal source. | (Drugman, Bozkurt, et al., 2012) |
| Cepstral Peak Slope (CPS) | Slope of the amplitude across all cepstral values was used as a measure of periodicity | (Skowronski et al., 2015) |
| Cepstral Peak Prominence (CPP) | Difference in amplitude between the peak cepstral value and the mean of all cepstral values was used as a measure of periodicity | (Fraile & Godino-Llorente, 2014) |
| Shape parameter of the Liljencrants-Fant Model of the Glottal Pulse Dynamic (RD) | Associated with dysphonic voice. | (Fant, 1997) |
| Creak | vocal glottal fry, reconstructed from multiple features | (Drugman et al., 2014; Drugman, Kane, et al., 2012; Ishi et al., 2008; Kane et al., 2013) |
| Clarity of articulation | relative depth of the minimum average magnitude difference function valley in the plausible pitch range | (Graciarena et al., 2013; Kotti & Stylianou, 2017; Ross et al., 1974; Samareh et al., 2018) |

#### S2. Informed and skeptical priors for differences by diagnostic group (methods)

Informed priors - when available - were normal distributions based on the meta-analytic effects (see table 2). Skeptical priors were modeled as normal distribution centered at zero (no group difference), with a standard deviation of .3. The expected estimates of the skeptical model were thus predominantly included within -.9 and .9, but could still be swayed by the data. Individual variability was modeled using a positive half-normal prior centered at 0, with a standard deviation of 0.1 from the estimate for the specific group and language of the individual, thus regularizing the inference.

#### S3 – Relations to clinical studies (methods)

To analyse the relation of clinical and acoustic features (ADOS total, Communication, Social Interaction, Repetitive Behaviors scores) we built multilevel Bayesian linear regression models with the acoustic feature as outcome and clinical features as ordinal predictors, on the autistic group only, separately by language and with varying effects by participant. While we had clinical scores for many of the neurotypical children as well, the ADOS scale is designed to assess autistic people, and therefore NT children present very minimal variability in scores, with a clearly distinct mode from scores in ASD. Including ADOS data from the neurotypical children would thus introduce severe bias in the models. We wanted to facilitate comparisons with the previous literature cited in the meta-analysis reporting Pearson correlation coefficients, however we could not simply standardize the clinical features as z-scores, as that would assume a linear relation between acoustic and clinical features. Clinical features are more adequately modeled as ordinal variables, where a monotonic relation is assumed but non-linear forms are possible. In other words, while we expect that if there is a change in acoustic patterns when moving from an ADOS Communication score of 0 to a score of 1, we should see a change in the same direction when moving from 1 to 2, but the size of the change might be different. We therefore opted to rescale the acoustic features on a 0-1 scale, maintain the clinical features as monotonic, and report the posterior estimates of the relation between clinical and acoustic features on a Pearson r scale, that is, as the effects of a change from minimum to maximum clinical score. Note that in case of linear changes, the current model gives comparable estimates to more traditional models (Bürkner & Charpentier, 2020). Apart from this difference, we followed the procedure described in the previous paragraphs.

### S4. Interdependencies between features and feature selection

Expanding the acoustic features investigated will produce a non-trivial increase in the number of statistical analyses required, potentially inflating the risk of false positives. Further, acoustic features are likely to be related to each other, and therefore we should assess whether all the features investigated provide independent information, and whether it is really necessary to add more complex acoustic measures of voice quality to the more traditional prosodic measures. Broadly speaking, there are at least four main approaches to the problem of feature-space reduction: 1) theoretically-justified a priori decisions, 2) dimensionality reduction methods, 3) clustering techniques, and 4) outcome-based methods. Each of these is a potentially viable method for reducing the feature space, but each comes with trade-offs. We briefly discuss each of these in turn.

Theoretically-justified a priori feature selection is the simplest of these. Choosing features a priori has the advantage of being perhaps the most easily interpretable of all four approaches: given that the features have been chosen on theoretically informed grounds, the framework for interpreting them is already there.

Dimensionality reduction methods, such as Principal Component Analysis (PCA) are a class of methods which involve the data-driven inference of latent variables underlying the actual features investigated. The goal is to identify a small number of variables which can account for the majority of the variance in the acoustic features (Pearson, 1901). Given the large number of acoustic features to choose from, PCA is commonly used to identify a smaller number of inferred features and reduce the original complexity without losing the original information (e.g. in Cohen et al., 2016). However, one should be careful with PCA: since it does not distinguish between shared and unique variance among the original features, the components identified may be difficult to interpret within a theoretically meaningful framework (Preacher & MacCallum, 2003).

Network modeling approaches conceive of features as nodes in a network, and represent the shared variance between them graphically as connections between the nodes. While less commonly used, a key advantage to network models is that they represent the relationships between the original variables graphically, making them easier to interpret. A variety of algorithms exist for identifying “communities” of related variables, thus facilitating dimensionality reduction. Here we focus on spin-glass community detection algorithms, which are taken from the so-called Potts model in statistical physics, and seeks to identify sets of variables with as many positive correlations between community members as possible, and either as many negative correlations with variables outside the community as possible, or simply as few connections with nodes outside the group as possible (Reichardt & Bornholdt, 2006; Traag & Bruggeman, 2009). More generally, communities are defined as a set of variables that share more variance with each other than with variables not belonging to the community.

A final approach is outcome-driven (or supervised) feature selection. This common machine learning approach aims at identifying the minimum set of features most effective in discriminating between groups (Huang, 2015; Smialowski et al., 2010). These algorithms evaluate features based on their correlations with previously labelled data (Sheikhpour et al., 2017). Because this approach selects features from the original dataset, it can maintain a reasonable degree of interpretability, although these techniques can easily choose a combination of features that do not make obvious intuitive sense.

We wished to explore common principled means of reducing these often intercorrelated acoustic features to a smaller subset of features. Ideally, these should be features which are not only useful for modeling the voice in a predictive framework, but which are also easily generalizable and clinically intuitive. We therefore chose to focus on dimensionality reduction and network analysis. We discarded a priori feature selection, although we hope that over time, cumulative and theory-driven studies will lead to an a priori set of features. We also set aside outcome-based methods as our primary interest here is in understanding the broader landscape of acoustic features associated with the speech of people with autism, and not optimizing for predictive power with our particular dataset.

We apply principal component analysis (PCA) and a network-based spin-glass community detection algorithm separately to each dataset.

#### S5 - Implementation notes

Bayesian meta-analyses were calculated using the brms (Bürkner, 2017) R interface for Stan (Gelman et al., 2015). PCA’s were calculated and explored using the FactoMineR (Lê et al., 2008) and factoextra (Kassambara & Mundt, 2017) packages for R. Network modelling and community detection were done using the qgraph (Epskamp et al., 2012), igraph (Csardi & Nepusz, 2006), and ggraph (Pedersen, 2017) packages for R. Figures were produced with ggplot2 from the tidyverse package (Wickham & Wickham, 2017). All analyses were conducted in R using the RStudio IDE (R Core Team, 2018; RStudio Team, 2020).

#### S6 – Differences by diagnostic group – novel features (results)

Table S2. Estimated standardized mean differences (ASD – NT) for the novel acoustic measures. The first column reports the main effect of the diagnostic group (across sex and age), respectively for Danish and for US English. The second column indicates the interaction between the effect of diagnostic group and biological sex (Male – Female), that is, the difference in effect of group between the male and the female participants. The third column reports the interaction between the effect of diagnostic group and age, that is, the change in effect size as age increases of 1 standard deviation. ER indicates the evidence ratio for the difference, ER01 the evidence ratio for the null effect.

|  | Group (ASD - NT) | Biological sex (M - F) | Age |
| --- | --- | --- | --- |
| *Jitter* |  |  |  |
| DK | 0 (-0.19 0.2) ER = 0.98 ER01 = 2.48 | 0.13 (-0.21 0.46) ER = 2.67 ER01 = 1.7 | 0 (-0.05 0.04) ER = 1.2 ER01 = 10.88 |
| US | **-0.39 (-0.58 -0.19) ER > 1000** | **0.34 (-0.11 0.78) ER = 9.03** | 0 (-0.03 0.03) ER = 1 ER01 = 13.98 |
| *Shimmer* |  |  |  |
| DK | **0.25 (0.06 0.44) ER = 58.7** | **-0.16 (-0.49 0.16) ER = 3.88** | **0.04 (-0.01 0.08) ER = 11.7** |
| US | **-0.35 (-0.53 -0.17) ER = 570.43** | 0.05 (-0.37 0.46) ER = 1.44 ER01 = 1.66 | 0 (-0.03 0.03) ER = 1.42 ER01 = 14.85 |
| *NAQ Median* |  |  |  |
| DK | **0.11 (-0.07 0.27) ER = 5.71** | 0.02 (-0.28 0.3) ER = 1.2 ER01 = 2.42 | **-0.02 (-0.07 0.02) ER = 3.97** |
| US | **0.1 (-0.04 0.24) ER = 7.7** | -0.11 (-0.51 0.29) ER = 2.03 ER01 = 1.67 | -0.01 (-0.04 0.02) ER = 3.42 |
| *NAQ IQR* |  |  |  |
| DK | **-0.06 (-0.21 0.09) ER = 3.01** | **0.45 (0.16 0.73) ER = 284.71** | **-0.04 (-0.09 0) ER = 16.62** |
| US | **0.16 (-0.04 0.36) ER = 8.8** | -0.09 (-0.53 0.35) ER = 1.67 ER01 = 1.45 | **-0.02 (-0.05 0.02) ER = 3.66** |
| *QOQ Median* |  |  |  |
| DK | 0.05 (-0.12 0.22) ER = 2.11 ER01 = 2.59 | 0.1 (-0.19 0.4) ER = 2.52 ER01 = 1.99 | **-0.03 (-0.07 0.02) ER = 5.71** |
| US | **0.1 (-0.05 0.24) ER = 6.89** | -0.1 (-0.49 0.29) ER = 1.95 ER01 = 1.69 | **-0.01 (-0.04 0.02) ER = 3.5** |
| *QOQ IQR* |  |  |  |
| DK | 0 (-0.15 0.15) ER = 1.06 ER01 = 3.38 | **0.43 (0.14 0.7) ER = 141.86** | 0 (-0.05 0.04) ER = 1.29 ER01 = 11.33 |
| US | 0.15 (-0.03 0.34) ER = 9.84 | -0.1 (-0.56 0.35) ER = 1.77 ER01 = 1.52 | 0 (-0.04 0.03) ER = 1.18 ER01 = 13.6 |
| *H1H2 Median* |  |  |  |
| DK | **-0.1 (-0.28 0.08) ER = 4.65** | **-0.21 (-0.48 0.07) ER = 9.1** | **-0.04 (-0.08 0.01) ER = 11.58** |
| US | 0.01 (-0.21 0.24) ER = 1.19 ER01 = 2.3 | **0.33 (-0.03 0.7) ER = 14.62** | -0.02 (-0.05 0.02) ER = 4.3 |
| *H1H2 IQR* |  |  |  |
| DK | 0.02 (-0.11 0.16) ER = 1.52 ER01 = 3.3 | **0.25 (-0.01 0.51) ER = 17.35** | 0.01 (-0.03 0.06) ER = 2.59 ER01 = 10.14 |
| US | **0.11 (-0.07 0.28) ER = 5.49** | **-0.15 (-0.52 0.22) ER = 3.02** | 0 (-0.03 0.03) ER = 1.04 ER01 = 14.9 |
| *PSP Median* |  |  |  |
| DK | 0 (-0.15 0.15) ER = 1.06 ER01 = 3.23 | 0.06 (-0.24 0.35) ER = 1.62 ER01 = 2.18 | 0 (-0.04 0.04) ER = 1.2 ER01 = 11.94 |
| US | **0.13 (-0.02 0.28) ER = 13.23** | -0.23 (-0.66 0.2) ER = 4.46 | 0.02 (-0.01 0.05) ER = 4.16 |
| *PSP IQR* |  |  |  |
| DK | -0.06 (-0.22 0.09) ER = 2.8 ER01 = 2.61 | 0.1 (-0.21 0.41) ER = 2.38 ER01 = 1.92 | 0.02 (-0.02 0.06) ER = 3.09 |
| US | **0.09 (-0.08 0.25) ER = 4.03** | -0.15 (-0.59 0.31) ER = 2.52 ER01 = 1.37 | **0.01 (-0.02 0.05) ER = 3.23** |
| *HRF Median* |  |  |  |
| DK | 0.02 (-0.18 0.2) ER = 1.25 ER01 = 2.59 | **-0.15 (-0.49 0.18) ER = 3.29** | **0.03 (-0.02 0.07) ER = 5.31** |
| US | **-0.29 (-0.53 -0.06) ER = 47.19** | 0.06 (-0.31 0.44) ER = 1.58 ER01 = 1.79 | **0.02 (-0.02 0.06) ER = 4.55** |
| *HRF IQR* |  |  |  |
| DK | **0.08 (-0.06 0.22) ER = 4.88** | **0.19 (-0.1 0.48) ER = 5.8** | **0.01 (-0.03 0.05) ER = 1.93 ER01 = 10.49** |
| US | **-0.11 (-0.34 0.13) ER = 3.52** | -0.04 (-0.43 0.35) ER = 1.28 ER01 = 1.68 | 0.01 (-0.03 0.05) ER = 2 ER01 = 11.63 |
| *MDQ Median* |  |  |  |
| DK | **0.07 (-0.08 0.22) ER = 3.31** | **0.18 (-0.07 0.44) ER = 7.32** | **-0.02 (-0.06 0.02) ER = 3.57** |
| US | **0.06 (-0.05 0.17) ER = 4.59** | **-0.2 (-0.55 0.15) ER = 4.85** | **0.02 (-0.01 0.04) ER = 6.37** |
| *MDQ IQR* |  |  |  |
| DK | **-0.08 (-0.25 0.08) ER = 3.88** | **-0.24 (-0.53 0.06) ER = 9.93** | 0.01 (-0.04 0.05) ER = 1.43 ER01 = 11.38 |
| US | -0.03 (-0.16 0.11) ER = 1.66 ER01 = 3.83 | 0.07 (-0.32 0.46) ER = 1.6 ER01 = 1.8 | 0 (-0.03 0.03) ER = 1.05 ER01 = 17.9 |
| *CPS Median* |  |  |  |
| DK | **0.27 (0.11 0.44) ER = 249** | **0.46 (0.18 0.75) ER = 165.67** | 0 (-0.04 0.04) ER = 1.14 ER01 = 11.43 |
| US | **-0.11 (-0.25 0.02) ER = 11.46** | 0.09 (-0.25 0.42) ER = 2.02 ER01 = 1.84 | **0.04 (0.01 0.06) ER = 69.18** |
| *CPS IQR* |  |  |  |
| DK | 0.06 (-0.1 0.22) ER = 2.69 ER01 = 2.66 | **0.48 (0.2 0.74) ER = 499** | -0.01 (-0.05 0.03) ER = 1.52 ER01 = 11.47 |
| US | **-0.16 (-0.38 0.06) ER = 7.11** | **0.26 (-0.18 0.7) ER = 5.01** | 0 (-0.03 0.04) ER = 1.46 ER01 = 13.73 |
| *Rd Median* |  |  |  |
| DK | 0 (-0.19 0.18) ER = 1.01 ER01 = 2.86 | **0.46 (0.14 0.76) ER = 124** | -0.01 (-0.05 0.04) ER = 1.64 ER01 = 10.62 |
| US | 0.04 (-0.1 0.18) ER = 2.27 ER01 = 3.15 | 0.07 (-0.36 0.49) ER = 1.54 ER01 = 1.61 | 0 (-0.03 0.03) ER = 1.26 ER01 = 16.92 |
| *Rd IQR* |  |  |  |
| DK | **-0.11 (-0.3 0.07) ER = 5.01** | **0.36 (0.03 0.69) ER = 26.4** | 0 (-0.05 0.04) ER = 1.14 ER01 = 10.45 |
| US | 0 (-0.13 0.14) ER = 1.06 ER01 = 3.67 | 0.01 (-0.39 0.42) ER = 1.07 ER01 = 1.77 | 0 (-0.03 0.03) ER = 1.01 ER01 = 16.39 |
| *Harmonicity Median* |  |  |  |
| DK | **-0.38 (-0.56 -0.21) ER > 1000** | **0.24 (-0.1 0.59) ER = 7.08** | **-0.02 (-0.07 0.02) ER = 3.59** |
| US | **0.15 (-0.01 0.31) ER = 16.17** | -0.15 (-0.57 0.25) ER = 2.56 ER01 = 1.38 | -0.01 (-0.04 0.02) ER = 2 ER01 = 13.98 |
| *Harmonicity IQR* |  |  |  |
| DK | **-0.15 (-0.33 0.03) ER = 12.29** | **-0.31 (-0.66 0.05) ER = 11.86** | -0.01 (-0.05 0.04) ER = 1.43 ER01 = 10.57 |
| US | **0.13 (-0.08 0.34) ER = 5.63** | **0.32 (-0.11 0.75) ER = 7.95** | **0.02 (-0.02 0.06) ER = 4.38** |
| *Clarity Median* |  |  |  |
| DK | **-0.38 (-0.56 -0.21) ER > 1000** | **0.24 (-0.1 0.58) ER = 7.83** | **-0.02 (-0.06 0.02) ER = 3.37** |
| US | **0.15 (0 0.31) ER = 17.35** | -0.15 (-0.57 0.27) ER = 2.63 ER01 = 1.4 | -0.01 (-0.04 0.02) ER = 1.94 ER01 = 15.33 |
| *Clarity IQR* |  |  |  |
| DK | **-0.15 (-0.32 0.03) ER = 10.9** | **-0.3 (-0.64 0.04) ER = 12.65** | -0.01 (-0.05 0.04) ER = 1.58 ER01 = 10.72 |
| US | **0.13 (-0.09 0.34) ER = 5.22** | **0.31 (-0.14 0.76) ER = 6.68** | **0.02 (-0.01 0.05) ER = 4.41** |
| *CPP Median* |  |  |  |
| DK | -0.03 (-0.23 0.18) ER = 1.39 ER01 = 2.44 | 0.14 (-0.23 0.51) ER = 2.65 ER01 = 1.61 | 0.01 (-0.03 0.06) ER = 2.1 ER01 = 10.13 |
| US | **0.21 (0.01 0.41) ER = 21.6** | -0.18 (-0.6 0.25) ER = 2.98 ER01 = 1.3 | 0.01 (-0.02 0.05) ER = 2.98 ER01 = 11.27 |
| *CPP IQR* |  |  |  |
| DK | 0.08 (-0.11 0.28) ER = 2.84 ER01 = 1.96 | **0.38 (0.05 0.73) ER = 37.83** | 0 (-0.04 0.05) ER = 1.22 ER01 = 10.35 |
| US | **0.31 (0.11 0.51) ER = 124** | -0.01 (-0.48 0.45) ER = 1.06 ER01 = 1.51 | 0.01 (-0.02 0.05) ER = 3 ER01 = 11.62 |
| *Creak Median* |  |  |  |
| DK | **0.15 (0.07 0.23) ER = 399** | **0.11 (-0.07 0.28) ER = 5.22** | 0.01 (-0.03 0.04) ER = 1.74 ER01 = 14.26 |
| US | **0.13 (-0.13 0.39) ER = 4.21** | **-0.27 (-0.67 0.13) ER = 6.59** | **0.03 (-0.01 0.07) ER = 8.76** |
| *Creak IQR* |  |  |  |
| DK | **0.07 (-0.01 0.14) ER = 13.71** | 0.03 (-0.16 0.21) ER = 1.46 ER01 = 3.85 | 0 (-0.04 0.03) ER = 1.26 ER01 = 14.45 |
| US | -0.09 (-0.35 0.16) ER = 2.66 ER01 = 1.58 | 0.07 (-0.32 0.45) ER = 1.65 ER01 = 1.63 | 0.01 (-0.03 0.05) ER = 1.62 ER01 = 12.09 |

#### S7 – Dimensionality reduction (results)

Principal Component Analysis did not yield any insights into how the feature-space could be meaningfully reduced, see Figure S1. The first 10 principal components for each group cumulatively accounted for a substantial portion of the variance (DK NT = 87.8%, DK ASD = 87.7%, US NT = 93.5%, USA NT = 93.4%). However, the distribution of variance across the components was unequal between the two languages, with the first two components accounting for substantially more variance in the American English speakers than in the Danish speakers (see Figure S2). Features from the three feature types were fairly evenly distributed along components 1 and 2, and no clear patterns were discernable. Both the Kaiser-Guttman rule and the scree plot method indicated that the feature space of the data could be reduced to approximately 7-9 components (see Figure S3). Inspection of the relative contributions to the components did not suggest any clear pattern, although a full exploration of these data is beyond the scope of this paper.

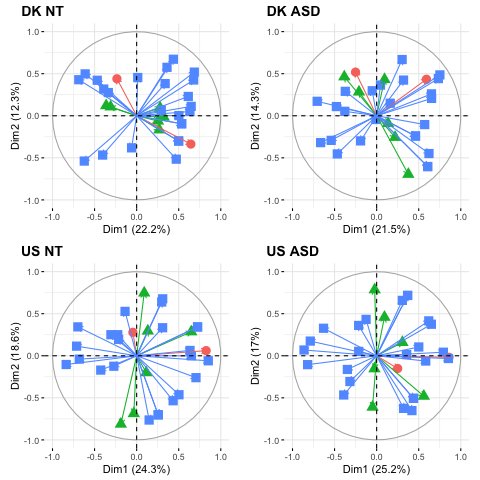

*Figure S1: Features plotted against the first two principal components (Dimension 1 and Dimension 2). Green triangles represent rhythm features, red circles represent pitch features, and blue square represent voice quality features.*

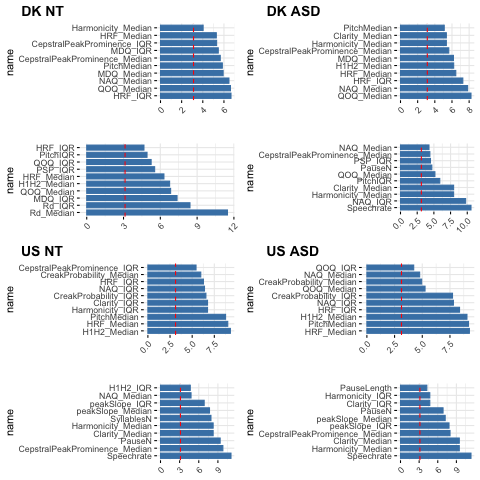

Figure S2: Feature contributions to the first and second principal components

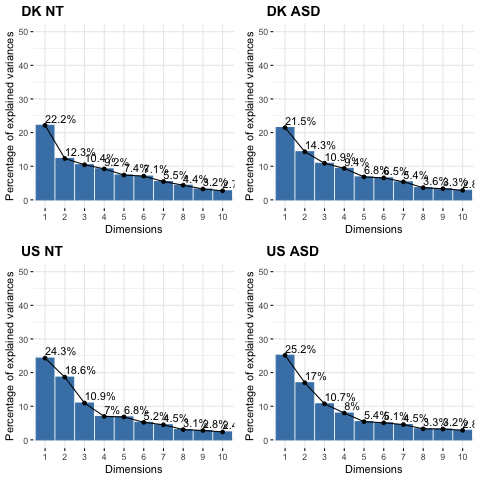

Figure S3: Scree plots of the first 10 components from the PCA analysis

Spin-glass community detection did not yield any immediate insights into how the feature-space could be meaningfully reduced, either, see Figure S4. Three of the four groups settled on a three-community solution, while the fourth (DK NT) settled on a two-community solution. There was some indication that underlying patterns may exist, e.g., articulation rate, speech rate, and number of syllables were clustered in the same community in all four groups (see Table S3), however a full exploration of these data is beyond the scope of this paper.

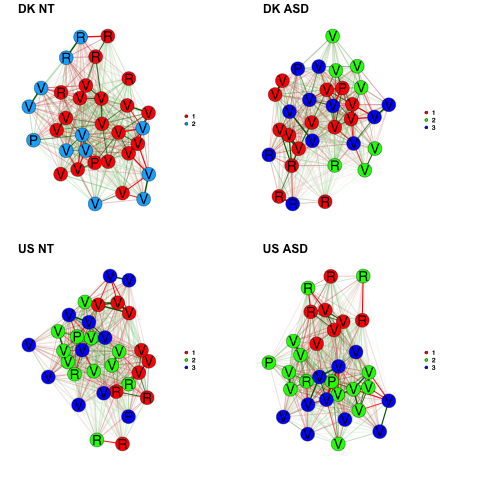

*Figure S4: Partial correlation network graphs of acoustic features. Letters indicate the type of acoustic feature: “P” pertains to pitch; “R” to rhythm and duration; “V” to voice quality. Colors indicate feature communities identified with spin-glass algorithm (Traag & Buggerman, 2009).*

*Table S3 – Communities identified by the spin-glass community detection algorithm. Communities are reported for each feature within each dataset separately and identified with a number (1-3)*

| DK NT | Com | DK ASD | Com | US NT | Com | US ASD | Com |
| --- | --- | --- | --- | --- | --- | --- | --- |
| ArticulationRate | 1 | ArticulationRate | 1 | ArticulationRate | 1 | ArticulationRate | 1 |
| PauseLength | 1 | Speechrate | 1 | Speechrate | 1 | Speechrate | 1 |
| Speechrate | 1 | SyllablesN | 1 | SyllablesN | 1 | SyllablesN | 1 |
| SyllablesN | 1 | PitchMedian | 1 | NAQ_Median | 1 | Harmonicity_Median | 1 |
| PitchMedian | 1 | NAQ_Median | 1 | NAQ_IQR | 1 | Clarity_Median | 1 |
| NAQ_Median | 1 | QOQ_Median | 1 | QOQ_Median | 1 | CepstralPeakProminence_Median | 1 |
| NAQ_IQR | 1 | H1H2_Median | 1 | Harmonicity_Median | 1 | CepstralPeakProminence_IQR | 1 |
| QOQ_Median | 1 | MDQ_Median | 1 | Clarity_Median | 1 | PauseLength | 2 |
| H1H2_Median | 1 | Harmonicity_Median | 1 | CepstralPeakProminence_Median | 1 | PauseN | 2 |
| MDQ_Median | 1 | Harmonicity_IQR | 1 | PauseLength | 2 | SyllableLength | 2 |
| peakSlope_Median | 1 | Clarity_Median | 1 | PauseN | 2 | PitchMedian | 2 |
| peakSlope_IQR | 1 | Clarity_IQR | 1 | SyllableLength | 2 | PitchIQR | 2 |
| Rd_Median | 1 | CepstralPeakProminence_Median | 1 | PitchMedian | 2 | NAQ_Median | 2 |
| Rd_IQR | 1 | CepstralPeakProminence_IQR | 1 | QOQ_IQR | 2 | NAQ_IQR | 2 |
| Harmonicity_Median | 1 | PauseLength | 2 | H1H2_Median | 2 | QOQ_Median | 2 |
| Harmonicity_IQR | 1 | NAQ_IQR | 2 | MDQ_Median | 2 | QOQ_IQR | 2 |
| Clarity_Median | 1 | QOQ_IQR | 2 | peakSlope_Median | 2 | H1H2_Median | 2 |
| Clarity_IQR | 1 | peakSlope_Median | 2 | peakSlope_IQR | 2 | MDQ_Median | 2 |
| CepstralPeakProminence_Median | 1 | peakSlope_IQR | 2 | Harmonicity_IQR | 2 | peakSlope_Median | 2 |
| CepstralPeakProminence_IQR | 1 | Rd_Median | 2 | Clarity_IQR | 2 | peakSlope_IQR | 2 |
| PauseN | 2 | Rd_IQR | 2 | CepstralPeakProminence_IQR | 2 | Harmonicity_IQR | 2 |
| SyllableLength | 2 | PauseN | 3 | PitchIQR | 3 | Clarity_IQR | 2 |
| PitchIQR | 2 | SyllableLength | 3 | H1H2_IQR | 3 | H1H2_IQR | 3 |
| QOQ_IQR | 2 | PitchIQR | 3 | PSP_Median | 3 | PSP_Median | 3 |
| H1H2_IQR | 2 | H1H2_IQR | 3 | PSP_IQR | 3 | PSP_IQR | 3 |
| PSP_Median | 2 | PSP_Median | 3 | HRF_Median | 3 | HRF_Median | 3 |
| PSP_IQR | 2 | PSP_IQR | 3 | HRF_IQR | 3 | HRF_IQR | 3 |
| HRF_Median | 2 | HRF_Median | 3 | MDQ_IQR | 3 | MDQ_IQR | 3 |
| HRF_IQR | 2 | HRF_IQR | 3 | Rd_Median | 3 | Rd_Median | 3 |
| MDQ_IQR | 2 | MDQ_IQR | 3 | Rd_IQR | 3 | Rd_IQR | 3 |
| CreakProbability_Median | 2 | CreakProbability_Median | 3 | CreakProbability_Median | 3 | CreakProbability_Median | 3 |
| CreakProbability_IQR | 2 | CreakProbability_IQR | 3 | CreakProbability_IQR | 3 | CreakProbability_IQR | 3 |

Moving from qualitative descriptions of the speech of autistic people to a more robust quantitative acoustic profile requires an openness to the many measures of prosody and voice quality available. However, building a meaningful and interpretable acoustic profile will ultimately require paring this large feature-set down to a smaller set of core features that are predictive of diagnosis and possibly symptom severity. In this paper, we have explored two approaches to feature-set reduction: Principal Component Analysis and network community-detection. Neither of these approaches yielded in clear insight into the problem of feature-space reduction. Had the PCA resulted in similar components for the four groups, e.g., with a heavy weighting of rhythm features on component 1 and a heavy weighting of pitch features on component 2, one might begin to focus more attention on these features for further investigation. Had the network analysis resulted in e.g., four-community solutions for each of the groups, with one group primarily populated with rhythm features, the second with pitch, and the final two with subsets of the voice quality features, then one might have a stronger basis for selecting perhaps a single representative feature of each of these communities. Although the speech of autistic people is qualitatively described using adjectives such as “harsh” or “robotic” that suggest a combination of both prosodic and voice quality features, our exploratory results do not give any strong indication of which of the many measures available should be selected for future research.

#### S8 - Analyses on segments of 6 seconds

In the analyses reported in the main manuscript we focused on acoustic analyses of the full recordings of each single trial independently of their length. However, it might be argued that confounds might arise from comparing audios of different lengths. E.g. if a group produced consistently longer recordings, measures of variability might be accordingly increased, simply due to the longer time-series analyzed. To obviate that potential critique we also systematically clipped all audio recordings in non-overlapping segments of 6 seconds, removing recordings that were too short and final segments that did not reach the full 6 seconds of length. This resulted in 3365 segments, 1738 produced by autistic participants (880 in Danish, 858 in US English) and 1627 by NT participants (1096 in Danish, 531 in US English). We then repeated all analyses on these segments.

##### S8.1 Acoustic features with meta-analytic results

The detailed results are reported in Table S4 and figure S5, which report for each feature the meta-analytic results (in other words, the informed priors), as well as the results for both the skeptical and the informed analyses, now performed on segments of 6 seconds, instead of the whole recording. The results are analogous to those reported on the full audio recordings - except for syllable length - and generally supported our hypotheses. We generally replicate meta-analytic findings across both data sets (H1). Autistic participants tend to use higher and more varied pitch, as well as fewer and longer pauses and longer syllable length. Again, only in US English we see evidence of slower speech rate. Again we see smaller effect sizes in our data than in the meta-analysis (H1a). Using informed meta-analytic priors yielded bigger results than using skeptic ones, however they were still smaller than in the meta-analysis. Interestingly only half of the informed models (pitch median, number and length of pauses) performed better than the skeptic ones: including information from previous studies often made our statistical inference more robust and able to generalize to new data (LOO based stacking weights for informed models of 1), except when the meta-analytic information was in direct contradiction of the skeptical estimate in at least one language. Finally, the analyses indicate evidence of biological sex and age affecting the difference between autistic and neurotypical populations, but, again, inconsistently so across languages (H3).

Table S4 - Estimated standardized mean differences (ASD – NT) for the six acoustic measures present in the meta-analysis. The first column reports the main effect of the diagnostic group (across sex and age), respectively from the meta-analysis (MA), for the skeptical Danish and US English models , and for the informed ones. The second column indicates the interaction between the effect of diagnostic group and biological sex (Male – Female), that is, the difference in effect of group between the male and the female participants. The third column reports the interaction between the effect of diagnostic group and age, that is, the change in effect size as age increases of 1 standard deviation. ER indicates the evidence ratio for the difference, ER01 the evidence ratio for the null effect. Emphasis (in bold) indicates findings with more than anecdotal evidence.

|  | Group (ASD - NT) | Group_*  Biological sex (M - F) | Group_*  Age |
| --- | --- | --- | --- |
| *Pitch Median* | Informed model weight = 1 |  |  |
| MA | 0.38 (0.16 0.59) | NA | NA |
| Skeptical DK | **0.1 (-0.1 0.29) ER = 3.87** | **0.29 (-0.04 0.63) ER = 11.94** | -0.01 (-0.06 0.03) ER = 2.38 ER01 = 9.46 |
| Skeptical US | **0.37 (0.15 0.61) ER = 306.69** | -0.03 (-0.37 0.31) ER = 1.27 ER01 = 2.1 | **-0.02 (-0.06 0.01) ER = 5.3** |
| Informed DK | **0.27 (0.13 0.42) ER > 1000** | NA | NA |
| Informed US | **0.43 (0.27 0.58) ER > 1000** | NA | NA |
| *Pitch Variability* | Informed model weight=0.37 |  |  |
| MA | 0.48 (0.26 0.7) | NA | NA |
| Skeptical DK | **0.23 (0.09 0.38) ER = 136.93** | **0.14 (-0.18 0.45) ER = 3.41** | 0 (-0.04 0.04) ER = 1.04 ER01 = 11.16 |
| Skeptical US | -0.06 (-0.25 0.13) ER = 2.48 ER01 = 2.23 | **-0.21 (-0.55 0.14) ER = 4.93** | **-0.02 (-0.06 0.01) ER = 7.58** |
| Informed DK | **0.36 (0.24 0.48) ER > 1000** | NA | NA |
| Informed US | **0.23 (0.09 0.38) ER = 399** | NA | NA |
| *Speech Rate* | Informed model weight = 0 |  |  |
| MA | 0.02 (-0.27 0.31) | NA | NA |
| Skeptical DK | 0.04 (-0.1 0.18) ER = 2.03 ER01 = 3.13 | **-0.16 (-0.45 0.13) ER = 4.42** | 0.01 (-0.04 0.05) ER = 1.37 ER01 = 11.26 |
| Skeptical US | **-0.18 (-0.31 -0.04) ER = 53.05** | **0.19 (-0.14 0.52) ER = 5.17** | **-0.03 (-0.06 0) ER = 27.78** |
| Informed DK | 0.03 (-0.09 0.16) ER = 2.17 ER01 = 1.76 | NA | NA |
| Informed US | **-0.14 (-0.26 -0.02) ER = 33.78** | NA | NA |
| *Syllable Length* | Informed model weight = 0 |  |  |
| MA | 0.06 (-0.63 0.76) | NA | NA |
| Skeptical DK | **0.06 (-0.07 0.18) ER = 3.39** | **0.26 (-0.01 0.54) ER = 16.02** | -0.01 (-0.04 0.04) ER = 1.45 ER01 = 12.19 |
| Skeptical US | **0.13 (0.02 0.25) ER = 41.55** | **-0.15 (-0.46 0.17) ER = 3.83** | **0.03 (0 0.05) ER = 28.63** |
| Informed DK | **0.07 (-0.06 0.2) ER = 3.92** | NA | NA |
| Informed US | **0.14 (0.03 0.25) ER = 50.95** | NA | NA |
| *Pause Number* | Informed model weight = 1 |  |  |
| MA | 0.4 (0.01 0.78) | NA | NA |
| Skeptical DK | **-0.11 (-0.22 0) ER = 17.35** | 0.08 (-0.15 0.32) ER = 2.29 ER01 = 2.51 | -0.01 (-0.05 0.03) ER = 2.68 ER01 = 10.42 |
| Skeptical US | **-0.19 (-0.4 0.01) ER = 15.81** | **0.59 (0.19 0.98) ER = 128.03** | **-0.02 (-0.06 0.01) ER = 6.33** |
| Informed DK | **-0.05 (-0.16 0.06) ER = 3.72** | NA | NA |
| Informed US | **-0.09 (-0.27 0.09) ER = 4.08** | NA | NA |
| *Pause Length* | Informed model weight = 1 |  |  |
| MA | 0.21 (-0.09 0.5) | NA | NA |
| Skeptical DK | **0.39 (0.22 0.56) ER > 1000** | **0.25 (-0.1 0.59) ER = 7.3** | 0.01 (-0.03 0.06) ER = 2.17 ER01 = 10.31 |
| Skeptical US | 0 (-0.11 0.12) ER = 1.06 ER01 = 4.03 | **-0.17 (-0.48 0.15) ER = 4.26** | 0.01 (-0.02 0.03) ER = 2.18 ER01 = 18.33 |
| Informed DK | **0.37 (0.22 0.5) ER > 1000** | NA | NA |
| Informed US | **0.04 (-0.06 0.15) ER = 3.21** | NA | NA |

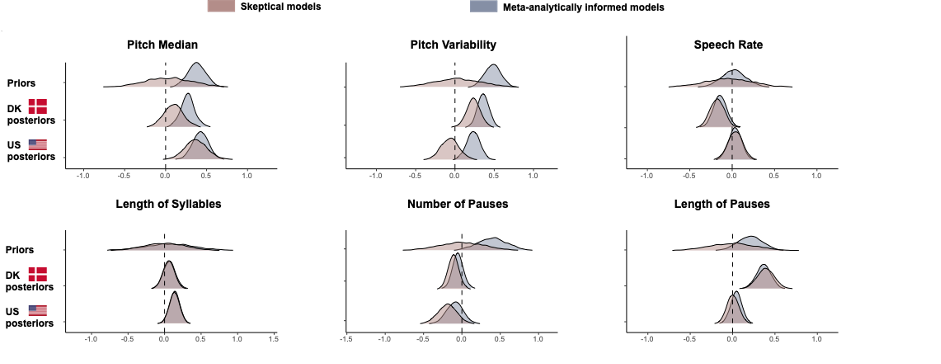

Figure S5: Results from informed and skeptic analyses across languages. Each panel represents the estimated differences by group (ASD - NT) in each language (demarcated by a flag), according to the use of skeptic (in red) or informed (in blue) priors.

##### S8.2 Novel acoustic features

The detailed results are reported in table S5. The results are similar, but not identical to the analysis of the full recordings. Generally we observe small to moderate (< 0.4) but reliable differences by group in the voice quality features within each data set, which are comparable to those in prosodic features (partially corroborating H2). As in more traditional acoustic features we see that including language, biological sex and age of the participants does in some cases affect the group differences (corroborating H3). Again, strikingly, only three acoustic measures (H1H2 IQR, MDQ and creak medians) present the same reliable relation with the diagnostic group across the two data sets (questioning the generalizability of H2).

Table S5 - Estimated standardized mean differences (ASD – NT) for the novel acoustic measures. The first column reports the main effect of the diagnostic group (across sex and age), respectively for Danish and for US English. The second column indicates the interaction between the effect of diagnostic group and biological sex (Male – Female), that is, the difference in effect of group between the male and the female participants. The third column reports the interaction between the effect of diagnostic group and age, that is, the change in effect size as age increases of 1 standard deviation. ER indicates the evidence ratio for the difference, ER01 the evidence ratio for the null effect. Emphasis (in bold) indicates findings with more than anecdotal evidence.

|  | Group (ASD - NT) | Group_*  Biological sex (M - F) | Group_*  Age |
| --- | --- | --- | --- |
| *Jitter* |  |  |  |
| DK | 0.02 (-0.16 0.2) ER = 1.31 ER01 = 2.6 | **0.16 (-0.17 0.51) ER = 3.67** | -0.01 (-0.05 0.04) ER = 1.62 ER01 = 10.71 |
| US | **-0.35 (-0.55 -0.16) ER = 499** | **0.38 (-0.06 0.81) ER = 13.34** | -0.01 (-0.04 0.02) ER = 2.28 ER01 = 13.06 |
| *Shimmer* |  |  |  |
| DK | **0.26 (0.07 0.45) ER = 104.26** | **-0.4 (-0.75 -0.07) ER = 37.83** | **0.04 (-0.01 0.09) ER = 14.75** |
| US | **-0.32 (-0.5 -0.15) ER = 799** | 0.11 (-0.29 0.51) ER = 2.03 ER01 = 1.7 | -0.01 (-0.04 0.02) ER = 1.96 ER01 = 16.22 |
| *NAQ Median* |  |  |  |
| DK | 0.07 (-0.09 0.24) ER = 2.91 ER01 = 2.68 | **0.14 (-0.15 0.44) ER = 3.78** | **-0.03 (-0.07 0.01) ER = 6.89** |
| US | **0.1 (-0.04 0.23) ER = 7.11** | -0.02 (-0.28 0.24) ER = 1.11 ER01 = 2.63 | **-0.02 (-0.05 0.01) ER = 5.47** |
| *NAQ IQR* |  |  |  |
| DK | **-0.06 (-0.2 0.08) ER = 3.24** | **0.26 (0.02 0.51) ER = 22.95** | **-0.04 (-0.08 0) ER = 13.98** |
| US | **0.16 (-0.04 0.35) ER = 10.3** | -0.02 (-0.37 0.32) ER = 1.18 ER01 = 2.04 | **-0.02 (-0.05 0.01) ER = 5.11** |
| *QOQ Median* |  |  |  |
| DK | -0.01 (-0.18 0.16) ER = 1.21 ER01 = 2.99 | **0.19 (-0.1 0.49) ER = 5.87** | **-0.04 (-0.08 0.01) ER = 11.16** |
| US | **0.1 (-0.05 0.24) ER = 6.48** | 0.01 (-0.25 0.27) ER = 1.13 ER01 = 2.52 | **-0.02 (-0.05 0.01) ER = 4.41** |
| *QOQ IQR* |  |  |  |
| DK | -0.01 (-0.16 0.15) ER = 1.16 ER01 = 3.34 | **0.12 (-0.14 0.36) ER = 3.62** | 0 (-0.04 0.04) ER = 1.04 ER01 = 12.29 |
| US | **0.1 (-0.09 0.28) ER = 4.35** | **-0.15 (-0.5 0.2) ER = 3.19** | 0 (-0.04 0.03) ER = 1.46 ER01 = 13.85 |
| *H1H2 Median* |  |  |  |
| DK | **-0.11 (-0.29 0.08) ER = 4.63** | -0.05 (-0.31 0.21) ER = 1.72 ER01 = 2.52 | **-0.04 (-0.08 0.01) ER = 12.47** |
| US | 0.06 (-0.15 0.26) ER = 2.18 ER01 = 2.06 | **0.37 (0.01 0.71) ER = 20.86** | **-0.02 (-0.05 0.02) ER = 4.05** |
| *H1H2 IQR* |  |  |  |
| DK | **0.06 (-0.06 0.18) ER = 4.18** | **0.19 (-0.04 0.42) ER = 10.98** | **0.02 (-0.02 0.05) ER = 3.03** |
| US | **0.07 (-0.09 0.24) ER = 3.25** | **-0.21 (-0.53 0.11) ER = 6.72** | 0 (-0.03 0.03) ER = 1.1 ER01 = 16.44 |
| *PSP Median* |  |  |  |
| DK | -0.05 (-0.2 0.1) ER = 2.27 ER01 = 2.64 | **0.12 (-0.16 0.4) ER = 3.1** | 0 (-0.05 0.04) ER = 1.33 ER01 = 11.01 |
| US | **0.15 (0.01 0.29) ER = 25.67** | **-0.18 (-0.49 0.12) ER = 5.37** | **0.02 (-0.01 0.05) ER = 4.2** |
| *PSP IQR* |  |  |  |
| DK | **-0.08 (-0.25 0.08) ER = 3.88** | 0.07 (-0.21 0.36) ER = 1.85 ER01 = 2.39 | **0.02 (-0.03 0.06) ER = 3.05** |
| US | **0.1 (-0.06 0.27) ER = 5.61** | **-0.17 (-0.49 0.16) ER = 4.4** | **0.02 (-0.02 0.05) ER = 4.04** |
| *HRF Median* |  |  |  |
| DK | 0.03 (-0.18 0.21) ER = 1.66 ER01 = 2.45 | **-0.21 (-0.54 0.12) ER = 5.64** | **0.03 (-0.02 0.07) ER = 4.52** |
| US | **-0.3 (-0.54 -0.08) ER = 53.79** | 0.02 (-0.3 0.35) ER = 1.11 ER01 = 2.1 | **0.02 (-0.02 0.06) ER = 4.75** |
| *HRF IQR* |  |  |  |
| DK | **0.1 (-0.03 0.22) ER = 9.81** | 0.04 (-0.24 0.31) ER = 1.43 ER01 = 2.36 | 0.01 (-0.03 0.05) ER = 2.25 ER01 = 11.51 |
| US | **-0.15 (-0.38 0.07) ER = 6.33** | -0.04 (-0.36 0.28) ER = 1.42 ER01 = 2.18 | 0.01 (-0.03 0.05) ER = 2.07 ER01 = 12.32 |
| *MDQ Median* |  |  |  |
| DK | **0.08 (-0.05 0.22) ER = 4.9** | **0.23 (-0.02 0.47) ER = 14.87** | **-0.02 (-0.07 0.02) ER = 4.49** |
| US | **0.08 (-0.02 0.18) ER = 8.98** | **-0.14 (-0.36 0.06) ER = 7.08** | **0.01 (-0.01 0.04) ER = 3.89** |
| *MDQ IQR* |  |  |  |
| DK | **-0.07 (-0.23 0.09) ER = 3.17** | **-0.34 (-0.62 -0.07) ER = 44.45** | 0.01 (-0.03 0.05) ER = 2.09 ER01 = 10.5 |
| US | -0.04 (-0.16 0.08) ER = 2.59 ER01 = 3.17 | 0.1 (-0.18 0.38) ER = 2.61 ER01 = 2.12 | 0 (-0.02 0.03) ER = 1.27 ER01 = 18.83 |
| *CPS Median* |  |  |  |
| DK | **0.27 (0.12 0.43) ER = 799** | **0.4 (0.12 0.66) ER = 99** | -0.01 (-0.05 0.03) ER = 1.61 ER01 = 11.34 |
| US | **-0.09 (-0.22 0.04) ER = 6.78** | 0.08 (-0.23 0.38) ER = 2.14 ER01 = 2.1 | **0.03 (0 0.06) ER = 33.78** |
| *CPS IQR* |  |  |  |
| DK | 0.01 (-0.12 0.14) ER = 1.23 ER01 = 3.56 | **0.45 (0.22 0.68) ER > 1000** | -0.01 (-0.06 0.03) ER = 2.6 ER01 = 10.11 |
| US | **-0.15 (-0.36 0.05) ER = 7.46** | **0.24 (-0.14 0.61) ER = 5.9** | 0 (-0.03 0.04) ER = 1.33 ER01 = 14.25 |
| *Rd Median* |  |  |  |
| DK | -0.01 (-0.2 0.19) ER = 1.08 ER01 = 2.51 | **0.66 (0.29 1.01) ER = 362.64** | 0 (-0.05 0.04) ER = 1.26 ER01 = 10.89 |
| US | **0.07 (-0.05 0.18) ER = 5.1** | **0.16 (-0.16 0.48) ER = 4.03** | 0 (-0.03 0.03) ER = 1.14 ER01 = 18.15 |
| *Rd IQR* |  |  |  |
| DK | **-0.15 (-0.34 0.04) ER = 9.34** | **0.56 (0.21 0.92) ER = 136.93** | -0.01 (-0.05 0.04) ER = 1.68 ER01 = 10.34 |
| US | 0.02 (-0.05 0.09) ER = 2.5 ER01 = 6.39 | 0.04 (-0.24 0.31) ER = 1.45 ER01 = 2.74 | -0.01 (-0.03 0.01) ER = 2.77 ER01 = 22.61 |
| *Harmonicity Median* |  |  |  |
| DK | **-0.32 (-0.47 -0.16) ER > 1000** | 0.12 (-0.19 0.43) ER = 2.81 ER01 = 1.88 | **-0.02 (-0.06 0.03) ER = 3.49** |
| US | **0.08 (-0.07 0.22) ER = 4.42** | **-0.14 (-0.47 0.18) ER = 3.24** | **-0.01 (-0.04 0.02) ER = 3.96** |
| *Harmonicity IQR* |  |  |  |
| DK | **-0.21 (-0.36 -0.06) ER = 74.47** | -0.03 (-0.34 0.3) ER = 1.28 ER01 = 2.2 | -0.02 (-0.06 0.03) ER = 2.59 ER01 = 9.9 |
| US | **0.1 (-0.09 0.3) ER = 4.07** | **0.21 (-0.18 0.62) ER = 4.07** | **0.02 (-0.02 0.05) ER = 3.73** |
| *Clarity Median* |  |  |  |
| DK | **-0.33 (-0.48 -0.17) ER > 1000** | **0.14 (-0.16 0.44) ER = 3.36** | **-0.02 (-0.06 0.02) ER = 3.16** |
| US | **0.08 (-0.07 0.23) ER = 4.4** | **-0.14 (-0.46 0.19) ER = 3.07** | **-0.01 (-0.04 0.02) ER = 3.36** |
| *Clarity IQR* |  |  |  |
| DK | **-0.21 (-0.35 -0.06) ER = 56.97** | -0.02 (-0.36 0.29) ER = 1.19 ER01 = 2.15 | -0.01 (-0.06 0.03) ER = 2.4 ER01 = 10.04 |
| US | **0.09 (-0.1 0.28) ER = 3.91** | **0.2 (-0.2 0.6) ER = 4.01** | **0.02 (-0.02 0.05) ER = 3.98** |
| *CPP Median* |  |  |  |
| DK | -0.03 (-0.22 0.15) ER = 1.6 ER01 = 2.48 | 0.08 (-0.26 0.41) ER = 1.8 ER01 = 1.9 | 0.01 (-0.04 0.05) ER = 1.56 ER01 = 10.09 |
| US | **0.23 (0.05 0.4) ER = 61.5** | **-0.16 (-0.51 0.18) ER = 3.41** | 0.01 (-0.03 0.04) ER = 1.7 ER01 = 13.73 |
| *CPP IQR* |  |  |  |
| DK | 0.08 (-0.13 0.27) ER = 2.77 ER01 = 2.02 | **0.33 (-0.02 0.68) ER = 14.33** | 0 (-0.05 0.05) ER = 1.01 ER01 = 10.67 |
| US | **0.27 (0.09 0.44) ER = 172.91** | -0.09 (-0.52 0.33) ER = 1.79 ER01 = 1.42 | 0 (-0.03 0.04) ER = 1.41 ER01 = 13.98 |
| *Creak Median* |  |  |  |
| DK | **0.06 (0 0.12) ER = 25.14** | **0.08 (-0.08 0.24) ER = 4.16** | -0.01 (-0.04 0.02) ER = 1.74 ER01 = 16.86 |
| US | 0.01 (-0.2 0.21) ER = 1.09 ER01 = 2.44 | -0.09 (-0.43 0.25) ER = 2.1 ER01 = 1.82 | **0.02 (-0.01 0.06) ER = 5.92** |
| *Creak IQR* |  |  |  |
| DK | **0.04 (-0.02 0.11) ER = 6.35** | -0.02 (-0.18 0.14) ER = 1.25 ER01 = 4.08 | 0 (-0.03 0.03) ER = 1.28 ER01 = 14.42 |
| US | **-0.14 (-0.38 0.1) ER = 4.66** | 0.11 (-0.22 0.44) ER = 2.28 ER01 = 1.87 | 0.01 (-0.03 0.05) ER = 2.47 ER01 = 10.93 |

#### S8.3 - Analysis of the relation between clinical and acoustic features

Detailed results are presented in table S6. Results are similar to those of the analysis of the full recordings, but generally weaker. Clinical features do correlate with acoustic features (corroborating H3) but most correlations are small (< 0.2 or 4% of the variance), and decreased in size compared to those in the full recordings (nothing > 0.4, that is , between 16% and 29% of the variance). We also note that ADOS total scores are generally related to acoustic features, and the effect size is most often of similar size as more specific subscales, indicating that acoustic features might not be more clearly related to specific clinical features than to general symptom severity. Finally, we highlight how the relations between acoustic and clinical features are rarely consistent across the two data sets, with the exception of speech rate.

Table S6 - Estimated standardized relation between acoustic and clinical features. ER indicates the evidence ratio for the difference, ER01 the evidence ratio for the null effect. Emphasis (in bold) indicates findings with more than anecdotal evidence.

|  | *ADOS Total*  β (95% CIs) | | *ADOS Communication*  β (95% CIs) | | | *ADOS Social*  β (95% CIs) | | | *ADOS Stereotyped*  β (95% CIs) |
| --- | --- | --- | --- | --- | --- | --- | --- | --- | --- |
| *Pitch Median DK* | 0.08 (-0.44 0.61) ER = 1.45 ER01 = 29.28 | | 0.08 (-0.23 0.4) ER = 2.22 ER01 = 12.75 | | | 0.08 (-0.35 0.55) ER = 1.65 ER01 = 16.85 | | | 0.07 (-1.15 1.43) ER = 1.14 ER01 = 4.04 |
| *Pitch Median US* | 0.11 (-0.21 0.44) ER = 2.35 ER01 = 39.89 | | 0.04 (-0.17 0.25) ER = 1.64 ER01 = 17.39 | | | **-0.16 (-0.39 0.08) ER = 6.42** | | | **0.1 (-0.09 0.3) ER = 3.94** |
| *Pitch IQR DK* | 0.06 (-0.16 0.26) ER = 2.26 ER01 = 66.12 | | **0.09 (-0.03 0.23) ER = 9.03** | | | 0.02 (-0.15 0.21) ER = 1.37 ER01 = 45.41 | | | -0.03 (-0.83 0.67) ER = 1.01 ER01 = 7.9 |
| *Pitch IQR US* | 0.03 (-0.15 0.21) ER = 1.61 ER01 = 80.19 | | -0.02 (-0.1 0.08) ER = 1.54 ER01 = 44.5 | | | 0.01 (-0.1 0.12) ER = 1.42 ER01 = 62.81 | | | 0 (-0.11 0.11) ER = 1.01 ER01 = 29.74 |
| *Speech Rate DK* | **-0.2 (-0.39 -0.06) ER = 107.11** | | **-0.16 (-0.32 -0.07) ER > 1000** | | | **-0.13 (-0.31 0) ER = 19.73** | | | **0.24 (-0.25 1.14) ER = 3.66** |
| *Speech Rate US* | **-0.06 (-0.17 0.05) ER = 4.22** | | **-0.03 (-0.08 0.02) ER = 5.77** | | | **-0.06 (-0.12 0) ER = 17.18** | | | -0.01 (-0.08 0.06) ER = 1.71 ER01 = 40.64 |
| *Pause Number DK* | 0.06 (-0.1 0.23) ER = 2.77 ER01 = 82.22 | | **0.04 (-0.06 0.16) ER = 3.34** | | | **0.06 (-0.08 0.2) ER = 3.04** | | | -0.2 (-1.05 0.33) ER = 2.58 ER01 = 8.47 |
| *Pause Number US* | -0.04 (-0.14 0.06) ER = 2.62 ER01 = 121.69 | | -0.02 (-0.08 0.04) ER = 2.67 ER01 = 56.67 | | | 0.01 (-0.05 0.08) ER = 1.34 ER01 = 105.79 | | | 0 (-0.06 0.06) ER = 1.1 ER01 = 50.62 |
| *Pause Length DK* | **0.2 (0.03 0.41) ER = 39** | | **0.08 (-0.03 0.22) ER = 6.45** | | | **0.22 (0.07 0.48) ER = 128.03** | | | **-0.38 (-1.43 0.12) ER = 7.66** |
| *Pause Length US* | **-0.08 (-0.25 0.09) ER = 3.69** | | **-0.04 (-0.13 0.06) ER = 3.26** | | | **-0.05 (-0.16 0.06) ER = 3.93** | | | -0.03 (-0.14 0.08) ER = 2.09 ER01 = 24.04 |
| *Jitter DK* | -0.01 (-0.36 0.33) ER = 1.02 ER01 = 44.29 | | **0.09 (-0.15 0.33) ER = 3.26** | | | -0.05 (-0.37 0.24) ER = 1.5 ER01 = 27.92 | | | -0.1 (-1.24 0.94) ER = 1.38 ER01 = 5.12 |
| *Jitter US* | **0.25 (0.16 0.34) ER > 1000** | | **0.07 (0.02 0.12) ER = 85.96** | | | **0.08 (0.01 0.17) ER = 44.98** | | | **0.17 (0.08 0.23) ER = 570.43** |
| *HNR Median DK* | **-0.36 (-0.72 -0.06) ER = 40.67** | | **-0.29 (-0.6 -0.1) ER = 180.82** | | | **-0.29 (-0.65 -0.03) ER = 26.59** | | | 0.31 (-0.6 1.59) ER = 2.72 ER01 = 3.91 |
| *HNR Median US* | -0.08 (-0.33 0.17) ER = 2.45 ER01 = 46.98 | | 0.01 (-0.1 0.12) ER = 1.42 ER01 = 34.73 | | | -0.05 (-0.18 0.07) ER = 2.78 ER01 = 45.83 | | | -0.04 (-0.24 0.11) ER = 1.79 ER01 = 16.56 |
| *HNR IQR DK* | 0.11 (-0.17 0.4) ER = 2.98 ER01 = 44.82 | | 0.04 (-0.15 0.27) ER = 1.91 ER01 = 22.07 | | | **0.13 (-0.11 0.4) ER = 4.35** | | | -0.26 (-1.31 0.49) ER = 2.81 ER01 = 5.26 |
| *HNR IQR US* | -0.08 (-0.31 0.13) ER = 2.8 ER01 = 53.33 | | -0.04 (-0.14 0.08) ER = 2.61 ER01 = 29.41 | | | **-0.18 (-0.3 -0.06) ER = 110.11** | | | **-0.09 (-0.21 0.03) ER = 9.03** |

#### S9 - Machine learning approaches to acoustic markers of ASD

In order to systematically build on the previous meta-analysis, in this study we focused on univariate approaches to acoustic atypicalities in ASD, that is, assessing whether there were differences between autistic and neurotypical participants, one acoustic feature at a time. An increasingly popular approach is to rely on machine learning techniques to identify the minimal combination of acoustic features that can reliably identify ASD in never-heard-before speech recordings. This approach seems particularly relevant for future studies. First, because the effect sizes we found were never exceedingly large, indicating poor predictive power towards diagnosis and clinical features; but also because our attempts at reducing the set of features based on their shared variance did not produce intuitive subsets of features. This suggests the need to combine several features to be able to better describe acoustic profiles in ASD.

Further, current approaches in the machine learning literature have adopted an alternative approach to the construction of acoustic features: instead of relying on traditionally meaningful acoustic features they rely more directly on the speech signal, for instance via MFCCs (Hansen et al., 2021; Sechidis et al., 2021) or even unsupervised learning of the features via convolutional neural networks(Schneider et al., 2019). These are all promising venues which we will further explore in future studies. However, current approaches to acoustic markers of ASD in the machine learning literature (see review in Fusaroli et al (2017)) suffer from many of the limitations in the univariate approaches: limited sample sizes with limited heterogeneity, lack of reproducibility and replication efforts. Machine learning techniques are not a solution to the need to build a cumulative yet self-correcting science of acoustic markers of ASD.

### S10 – Steps towards a cumulative yet self-critical approach

We summarize here the key steps in the way we operationalized a cumulative yet self-critical approach. We want to stress that this is not the only way to approach these issues, and not all these steps are necessary or always recommendable. Nevertheless, this summary might provide useful references to others attempting to follow our suggestions and improve upon them.

1. *Build where possible systematic reviews of the relevant literature*. Systematic reviews provide less biased overviews of the literature compared to narrative reviews based on what the researcher already knows or on unsystematic searches. Through the overview of the literature, it is possible to identify good practices, known confounds, and blindspots in the literature. In particular, in (Fusaroli et al., 2017) we attempted to provide explicit guidelines for future studies, and a discussion of each of the points and their implementation.
2. *Build where possible systematic meta-analyses of the relevant effects.* Meta-analyses are often biased and suffer from the “garbage-in-garbage-out” problem. However, they provide useful reference points as to effects to replicate, heterogeneity to expect, and can be operationalized in terms of meta-analytically informed priors. In particular, in (Fusaroli et al., 2017, 2018), besides detailed estimates, we openly released data (on metavoice.au.dk) and scripts, to fully successive studies to freely rely upon and improve the meta-analyses.
3. *Explicitly build new studies on systematic reviews.* When designing a new study, it is useful to explicitly take into account the recommendations developed in the systematic review. Sometimes the recommendations cannot be followed due to other constraints (e.g., economic and temporal constraints) or because the authors do not agree with them (self-criticality). That is not a problem, but it is best explicitly discussed, so that future studies can also build upon that discussion.
4. *Explicitly build new statistical analyses on systematic meta-analysis findings.* Meta-analytic findings about effects, but also heterogeneity of effects, can be explicitly included in new analyses in the form of informed priors (or “posterior passing”). Informed priors are best not used uncritically, since their use carries assumptions: e.g., that the populations are comparable and the heterogeneity of the samples not underestimated. Therefore, quality checks of the model fit should always be performed (see (Gelman et al., 2020)), for instance, assessing which outcomes the meta-analytic priors would imply for the current study, or overlaying the meta-analytic priors and the consequent posteriors to assess for undue influences. Further, a comparative approach – akin to prior sensitivity testing – should be implemented. The inferences relying on informed priors can be compared with others relying on skeptical priors, or systematically varying the confidence of the informed priors. Such comparison can be assessed in terms of robustness (within which boundaries do the results stay qualitatively the same) or generalizability (which priors lead to lower out-of-sample error). The comparative assessments should of course also be critically assessed, as they also rely on assumptions. E.g., that the current data are representative of data produced in new studies.
5. *Practice open science.* Beyond being transparent with design and analysis choices and their motivation, release your detailed estimates, scripts and data. Data cannot always be released in its raw format. However, anonymized pre-processed data as they enter the analysis can often be shared. Alternatively, there are ongoing attempts at creating repositories compliant with data privacy issues (White et al., n.d.; Williams et al., 2021), which could be used as an inspiration.
